# Specific *BRCA* and immune configurations determine optimal response to platinum-based chemotherapy in triple negative breast and ovarian carcinomas

**DOI:** 10.1101/2021.08.19.456799

**Authors:** Francesca Menghi, Kalyan Banda, Pooja Kumar, Robert Straub, Lacey Dobrolecki, Isabel V. Rodriguez, Susan E. Yost, Harshpreet Chandok, Marc R. Radke, Angela S. Zhu, George Somlo, Yuan Yuan, Michael T. Lewis, Elizabeth M. Swisher, Edison T. Liu

## Abstract

Loss of homologous recombination repair (HRR) via germline and somatic *BRCA1* or *BRCA2* gene mutations and via *BRCA1* promoter methylation has been associated with better response to platinum agents and PARP inhibitors, in both triple negative breast cancer (TNBC) and ovarian carcinoma (OvCa). A major conundrum arising from recent clinical studies is why cancers with *BRCA1* promoter methylation (*BRCA1*meth) respond more poorly as compared to those bearing mutations in *BRCA1* and *BRCA2* (*BRCA*mut), given the biologically equivalent HRR deficiency in both states. We dissected this problem through detailed genomic analyses of primary TNBC and OvCa cohorts, as well as experimentation with patient-derived xenograft (PDX) models and genetically engineered cell lines. Using the precise genomic scar of the tandem duplicator phenotype as a precise genomic indicator of BRCA1 deficiency, we found that, in all cohorts, *BRCA1*mut and *BRCA1*meth cancers share an equivalent degree of BRCA1-linked genomic rearrangements. Nonetheless, we consistently found that patients with *BRCA*mut cancers, but not those with *BRCA1*meth cancers, had significantly better response outcomes when compared to those with *BRCA* proficient cancers. When fully promoter methylated *BRCA1* PDX TNBCs were exposed to a single short course of platinum chemotherapy an unmethylated *BRCA1* promoter allele emerged in resultant tumors associated with an increase in *BRCA1* expression. A separate analysis of PDXs derived from treatment naïve TNBCs featured complete methylation of the *BRCA1* promoter, whereas those derived from post-chemotherapy TNBCs invariably had only partial methylation. PDXs with partial methylation were significantly associated with lower response rates to *in vivo* platinum-based therapy compared to those with complete promoter methylation. Using single cell clonal expansions from a partially *BRCA1*meth PDX, we confirmed that the reduced level of methylation was due to the demethylation of one of the *BRCA1* promoter alleles and not to the outgrowth of a non-methylated clone. Clinically, analysis of primary OvCas confirmed that high levels of *BRCA1* methylation were significantly associated with reduced *BRCA1* gene expression whereas cancers with lower levels of *BRCA1* methylation had expression levels approaching those found in *BRCA1* proficient cancers. These data suggest that unlike *BRCA*mut cancers, where HRR deficiency is achieved via mutations that are genetically ‘fixed’, *BRCA1*meth cancers are highly adaptive to genotoxin exposure and more likely to recover *BRCA1* expression, which may explain their poorer therapeutic response. We further found that an increased immune transcriptional signal, especially an elevated M1 macrophage signature, is associated with enhanced response to platinum-based chemotherapy only in patients with *BRCA* proficient cancers, in both TNBC and OvCa cohorts underscoring the importance of characterizing molecular heterogeneity to enhance predictive precision in assigning response probabilities in TNBC and OvCa.

## INTRODUCTION

We previously described how loss of BRCA1 activity, through either disruptive mutations or promoter methylation, results in a distinct genomic profile, which can manifest in one of three variations (i.e., tandem duplicator phenotype (TDP) group 1, TDP group 1/2 mix and TDP group 1/3 mix) hereby collectively named ‘TDP type 1’, comprising many tens to hundreds of isolated tandem duplications evenly distributed throughout the genome with a characteristic span size of 11 kb (Menghi et al., 2018). TDP type 1 is highly prevalent in both TNBC and ovarian carcinomas (OvCas), reflecting the common high frequency of BRCA1 abrogation in these two cancer types and raising the clinical impact of this form of genomic instability. The TDP-related genomic scars are identical in scale and distribution in both cancer types, and their association with BRCA1 deficiency is near absolute, suggesting that TNBC and OvCas are closely related in large part via the downstream genomic effects of BRCA1 deficiency (Cancer Genome Atlas Research et al., 2013). Recent evidence that homologous recombination deficiency (HRD) caused by loss of BRCA1 or BRCA2 (BRCA) activity determines high sensitivity to DNA inter-strand crosslinking agents, such as platinum-based chemotherapies, has spurred several studies assessing the value of *BRCA* status or BRCA-linked genomic scars as predictors of treatment response in both TNBC and OvCa, with conflicting outcomes (Abkevich et al., 2012; Stronach et al., 2018; Telli et al., 2016; Watkins et al., 2014). These conflicting outcomes may be the result of different methods to ascertain of BRCA deficiency, different chemotherapeutic regimens, different cancer types (TNBC *vs* OvCa), and disease states (primary *vs* metastatic disease). Since BRCA2-linked cancers never develop the TDP type 1 phenotype, TDP is currently the most precise measure for the specific form of HRD that results from BRCA1 deficiency. Therefore, TDP assessment provides a unique genomic forensic tool to ascertain critical exposure to BRCA1-specific deficiency in the development of TNBC and OvCa. Herein, we sought to use this precise functional measure of BRCA1 deficiency to reexamine genomic clinical data across primary TNBC and OvCa clinical cohorts, whose treatment regimen comprising a combination of platinum- and taxane-based chemotherapy provided a consistent basis for comparison. We confirmed our observations in PDX and engineered cell line models of TNBC. Our results show that tumors with disruptive mutations in either *BRCA1* or *BRCA2* (*BRCA*mut), those with *BRCA1* silencing through promoter methylation (*BRCA1*meth), and those that are proficient for both *BRCA1* and *BRCA2* (i.e., wild type for both *BRCA1* and *BRCA2* and without *BRCA1* promoter methylation, hereby called non*BRCA*) are distinct clinical-therapeutic entities in their response to platinum chemotherapies. The mechanistic heterogeneity of response to DNA damaging chemotherapy across the different *BRCA* states can explain many of the inconsistencies seen in clinical trials and provides an approach to formulate more precise predictive markers for therapeutic response.

## RESULTS

### *BRCA1* mutation and methylation result in the same genomic scars but associate with different therapeutic response in TNBC

To test the hypothesis that TDP type 1 status may be predictive of optimal response to platinum-based therapy, we assessed TDP status in a cohort of 42 patients with early stage TNBC, from the City Of Hope National Medical Center (COH TNBC cohort (Yuan et al., 2021), Figure 1A **and Tables S1-2**). All cancer specimens were collected as pre-treatment in a phase II neoadjuvant trial of carboplatin and NAB-paclitaxel. Consistent with our previous observation, 17 of the 19 TDP tumors were classified as TDP type 1 and were strongly associated with *BRCA1* mutation or promoter methylation (16/17 (94.1%) vs. 2/25 (8%) of the other TDP groups, *i.e.*, TDP groups 2, 2/3 mix and non TDP; P = 3.9E-5, Figure 1B). However, there was no association between TDP type 1 status and pathological complete response (pCR, Figure 1C and **Table S3**), which is commonly considered as the measure of optimal response in TNBC patient cohorts (Huang et al., 2020). The rates of pCR were higher among patients with *BRCA1* mutant cancers (*BRCA1*mut, 6/7, 85.7%) compared to those with either *BRCA1* proficient (which we call non*BRCA*, 11/24, 45.8%) or *BRCA1*meth (4/11, 36.4%) cancers, although the difference did not reach statistical significance (Figure 1D). Both patients with *BRCA2* mutant cancers achieved pCR (Figure 1A). Since breast cancers from both *BRCA1* and *BRCA2* germline mutation carriers have been previously shown to be sensitive to platinum-based chemotherapeutic regimens (Tutt et al., 2018), we grouped TNBCs harboring *BRCA1* and *BRCA2* disruptive mutations together (i.e., *BRCA*mut). When analyzed in this manner, patients with *BRCA*mut cancers (but not *BRCA1*meth cancers) were statistically more likely to achieve pCR than those with non*BRCA* cancers (88.9% vs. 40.9%, P = 0.033, Figure 1E). To exclude that the lower pCR rates associated with *BRCA1* methylation could be due to ineffective suppression of *BRCA1* expression, we compared the expression levels of *BRCA1*mut vs. *BRCA1*meth TNBCs as assessed by RNAseq. *BRCA1* expression was lowest in *BRCA1*meth cancers compared to both *BRCA1*mut and *BRCA1* proficient cancers (Figure 1F). In addition, *BRCA1*mut and *BRCA1*meth TNBCs shared similar tandem duplication numbers and span size distribution profiles (Figures 1G and 1H), implying equivalent loss of BRCA1 activity across the two modes of functional abrogation. These results also suggest that the BRCA1 effect on genomic instability in the form of TDP type 1 induction is separable to its effect on therapeutic response: a higher pCR rate was restricted to patients with *BRCA1*mut cancers, and the TDP status alone was not a determinant of chemotherapeutic response.

**Figure 1.**
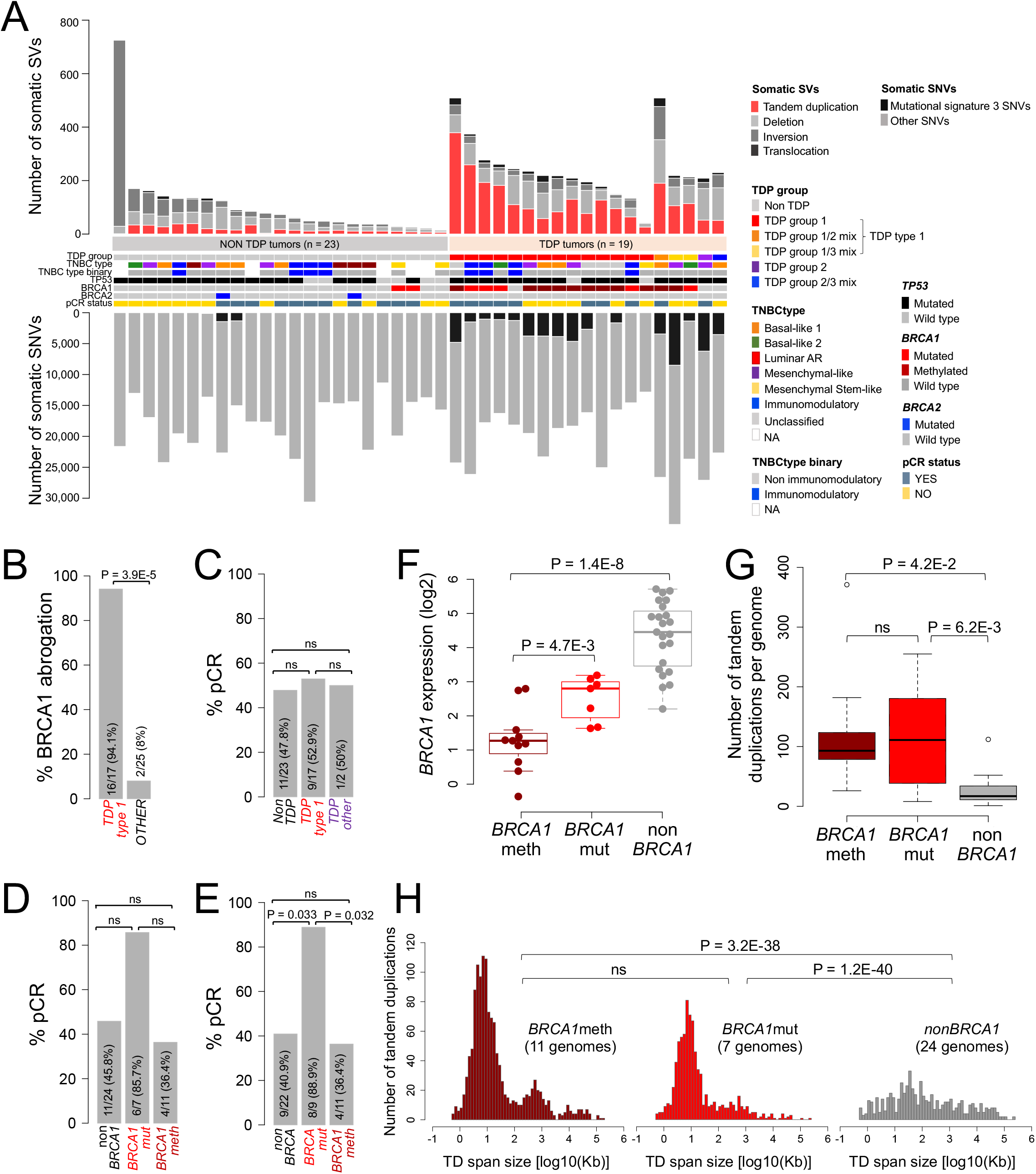
Genomic features and therapeutic response in the COH TNBC cohort. **A)** Overview of the TNBC cohort genomic and clinical features. Samples are sorted based on their TDP group assignment. **B)** Association between BRCA1 abrogation (by mutation or promoter methylation) and TDP status (TDP type 1 comprises all TDP subgroups with short span tandem duplications, i.e., TDP groups 1, 1/2 mix and 1/3 mix); P by logistic regression. **C-E)** Associations between TDP status (**C**, ‘TDP other’ refers to any TDP group other than TDP type 1) and *BRCA* mutation status (**D-E**) with pCR rates; P by logistic regression. **F)** *BRCA1* expression levels in *BRCA1*meth vs. *BRCA1*mut or non*BRCA1* TNBCs; P by Student’s t-test, two-tailed. **G)** Number of tandem duplications across *BRCA1*meth, *BRCA1*mut and non*BRCA1* cancer genomes; P by Student’s t-test, two-tailed. **H)** Tandem duplication span size distributions across *BRCA1*meth, *BRCA1*mut and non*BRCA1* TNBC genomes; P by Mann-Whitney test. Box plot elements: center line, median; box limits, lower and upper quartiles; whiskers extend up to one and a half times the interquartile range; ns, not significant.

### *BRCA* abrogation and specific response to carboplatin across PDX models of human TNBC

One of the objectives of our study was to ask whether TDP and/or *BRCA* status are the key determinants of platinum sensitivity in TNBC. However, most clinical studies (including those investigated herein) use combination chemotherapy which makes it difficult to parse out the effects of individual agents. To resolve this, we analyzed a cohort of 33 TNBC patient-derived xenografts (PDXs) with known response to carboplatin from an ongoing PDX preclinical trial at Baylor College of Medicine (https://pdxportal.research.bcm.edu/, to be described in detail elsewhere). Briefly, tumors were treated *in vivo* with single agent carboplatin (50 mg/kg, IP, weekly for four weeks), or single agent docetaxel (20 mg/kg, IP, weekly for four weeks). The PDX cohort comprised 11 *BRCA*mut PDXs carrying either *BRCA1* (n = 8) or *BRCA2* (n = 3) deleterious mutations, 6 *BRCA1*meth PDXs and 16 non*BRCA* PDXs (Figure 2A, **Tables S1-2**). As expected, we once more confirmed the strong association between BRCA1 abrogation and TDP type 1 (P < 0.001, Figure 2B). After a 28-day carboplatin regimen, we compared *in vivo* tumor growth rates between the different PDXs classified based on their *BRCA* status. We found that only 31.3% (5/16) of non*BRCA* PDX models treated with carboplatin showed a significant therapeutic response defined as a > 5% average decrease in tumor growth rate compared to the control arm, a highly quantitative response metric that is particularly useful when comparing across PDX studies (Hather et al., 2014). By contrast, 90.9% of *BRCA*mut PDXs (10/11, P = 0.009) had a response. *BRCA1*meth PDXs showed an intermediate response trend with 66.7% (4/6) of responders, although this was not statistically significant when compared to the non*BRCA* PDX models (Figure 2C, **Table S3**). Similar response profiles were obtained when response was assessed using modified RECIST criteria, where PDXs that demonstrated RECIST values equivalent to complete or partial response were designated as responders (Figures S1A-C and **Tables S2**). Importantly, no significant differences in response among the three *BRCA* classifications were observed when the same PDXs were treated with docetaxel as single agent (Figure 2D), validating our original hypothesis that the *BRCA* mutational status specifically sensitizes to platinum-based chemotherapy. Though TDP type 1 TNBC PDX models showed a trend towards better response to carboplatin, this was not statistically significant (Figure 2E, **Table S3**).

**Figure 2.**
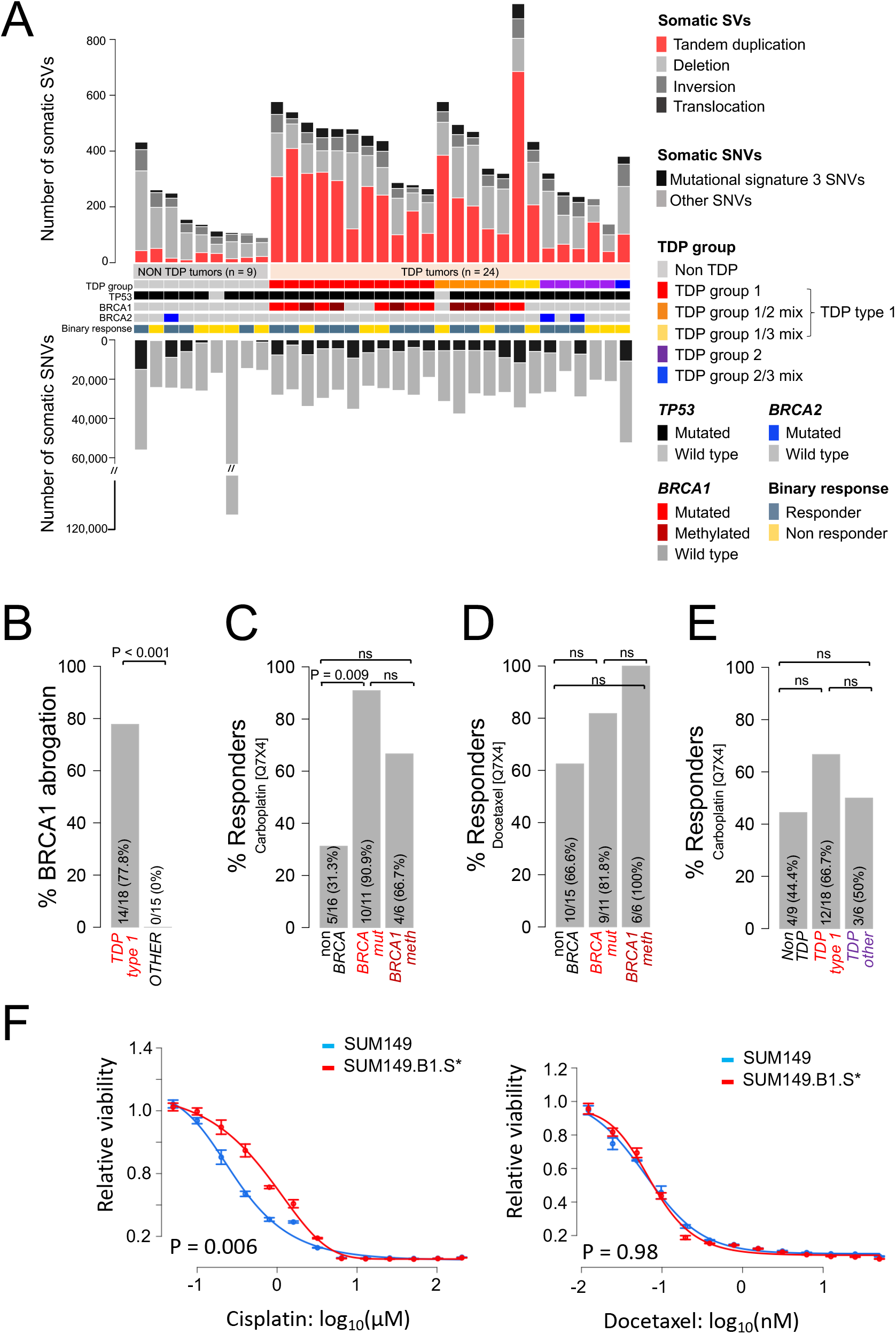
Genomic features and chemotherapy response in TNBC PDXs and cell lines. **A)** Overview of genomic features and chemotherapy response in the TNBC PDX cohort. Samples are sorted based on their TDP group assignment. **B)** Association between BRCA1 abrogation and TDP status; P by logistic regression; ns, not significant. **C-D)** Percentage of carboplatin (**C**) and docetaxel (**D**) responders as a function of *BRCA* status. P by logistic regression. **E)** Percentage of carboplatin responders as a function of TDP status in the PDX cohort. P by logistic regression. **F)** Cisplatin and docetaxel IC_50_ curves relative to the SUM149/SUM149.B1.S* isogenic cell lines. One representative example of four biological replicate experiments is shown in each graph. Data are presented as mean values and standard errors of the technical replicates. Significance values were calculated using the Student’s t-test (two-tailed) to compare IC_50_ values from the four biological replicates across the two cell lines.

To further confirm that the enhanced response in *BRCA*mut cancers is specific to *BRCA* gene mutations and not to other potentially associated mutations, we explored the chemotherapy responsiveness of the SUM149 TNBC cell line that harbors conjoint disruptive mutations in *BRCA1* and *TP53* as compared to its derivative SUM149.B1.S* cell line where BRCA1 activity was restored by introducing a secondary mutation via CRISPR-Cas9-mutagenesis, which reinstates the correct open reading frame (Drean et al., 2017). Our results showed that the SUM149 parental cell line was 2.6 times more sensitive to cisplatin (IC_50_ = 249 nM vs. 743 nM for the SUM149.B1.S* line; P = 0.006, Figure 2F, **Table S4**). Commonly, two other types of DNA damaging chemotherapeutic agents, doxorubicin and cyclophosphamide, are also used in combination for the treatment of TNBC. We found that the *BRCA1* mutant SUM149 parental line was 1.6 times more sensitive to doxorubicin and 6.2 times more sensitive to masfosfamide (a cyclophosphamide analog that does not require hepatic metabolism to be converted into its active form and it is therefore amenable to *in vitro* experimentation) than its BRCA1 proficient SUM149.B1.S* counterpart (Figure S1D, **Table S4)**. By contrast, the two cell lines did not differ in their IC_50_ values when treated with docetaxel, a taxane whose antitumor activity involves stabilization of cellular microtubules (IC_50_ = 101 pM for both cell lines, Figure 2F, **Table S4)**. This experiment confirms not only that mutations in the *BRCA1* gene are the cause of the differential sensitivity to platinum agents, but that this sensitivity is found for other DNA damaging agents but not for chemotherapies that target other cellular mechanisms, such as taxanes.

### Loss of *BRCA1* promoter methylation and induction of *BRCA1* expression in TNBC PDXs after platinum-based chemotherapy

We were puzzled as to why patients with *BRCA1*meth TNBCs did not show the same sensitivity to platinum-based chemotherapies compared to those with *BRCA1*mut TNBCs, given the functional equivalence between the two states in terms of the precise downstream genomic effects. To address this question, we first examined a total of 16 PDX models of *BRCA1*meth TNBC, combining the subset of *BRCA1*meth TNBCs from the PDX cohort described above and additional *BRCA1*meth TNBCs available from The Jackson Laboratory PDX Resource, all of which shared the TDP Type 1 configuration (**Table S5**). We observed two modes of *BRCA1* promoter methylation, as assessed by Methylation-Specific PCR (MSP): (a) complete methylation (i.e., no signal for the unmethylated PCR product) and (b) partial methylation (i.e., two signals corresponding to both the methylated and the unmethylated PCR products, Figure 3A). While MSP is not a quantitative assay *per se*, the absence of normal human DNA contamination in the PDX system justifies the assumption that a complete methylation profile corresponds to homozygous *BRCA1* methylation (i.e., all copies of the *BRCA1* promoter are methylated), whereas partial methylation indicates either a heterozygous profile or subclonal heterogeneity (Kondrashova et al., 2018). We then asked whether these different modes of promoter methylation had an impact on *BRCA1* gene expression. We found that complete promoter methylation associated with a 99% decrease in the median levels of *BRCA1* expression when compared to that of non*BRCA1* TNBC PDXs (p = 4.7E-07, Figure 3B), while partially *BRCA1*meth PDXs showed only a marginal reduction, with median *BRCA1* expression values reduced by only 41% (not significant, Figure 3B). Intriguingly, we also observed that while the great majority of the PDX models with complete methylation of the *BRCA1* promoter were established from treatment-naïve patient cancers (n = 9/11 (81.8%)), those which had been established from patient cancers after the exposure to chemotherapeutic regimens (i.e., post-treatment) consistently showed only partial methylation (n = 5/5, P=0.005, Figure 3C).

**Figure 3.**
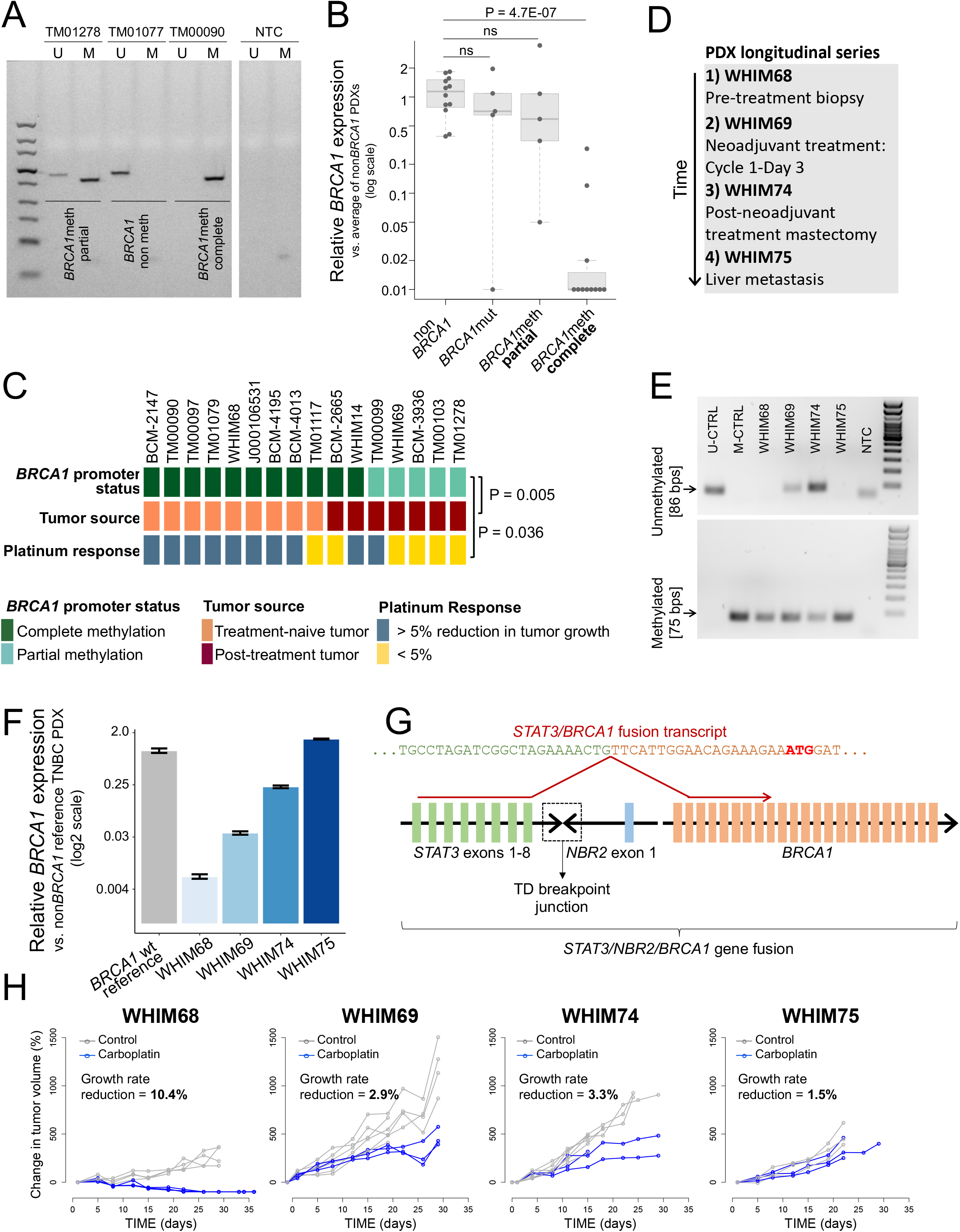
*BRCA1* methylation profiles and chemotherapy response in TNBC PDXs. **A)** MSP results for three exemplary TNBC PDX models, showing three different *BRCA1* methylation patterns at the proximal promoter (i.e., complete and partial methylation, and absence of methylation). U, unmethylated PCR product, M, methylated PCR product. **B)** *BRCA1* gene expression (qPCR) as a function of *BRCA1* status in TNBC PDX models. The average value of all the assessed non*BRCA1* PDX models was used as the calibrator in assessing expression fold changes. P values by Student’s t-test (two-tailed). **C)** *BRCA1* methylation status, tumor source and platinum-based therapy response for 16 *BRCA1*meth TNBC PDX models. Identical outcomes were obtained when response was determined using RECIST criteria, as reported in **Table S5**. P value by Fisher’s Exact test. **D)** Tumor source for the four TNBC PDX models established from the same patient donor (i.e., WHIM PDX longitudinal series). **F)** *BRCA1* expression levels for the four PDX models of the WHIM PDX longitudinal series, assessed by qPCR. A non*BRCA1* TNBC PDX was used as the calibrator. Data are presented as mean values and standard errors of the technical replicates. **G)** Schematic representation of the genomic fusion and corresponding fusion transcript involving the *STAT3*, *NBR2* and *BRCA1* genes in the post-treatment TNBC metastasis derived WHIM75 PDX model. Colored blocks represent individual exons and are not drawn to scale. A dashed box indicates the location of the tandem duplication breakpoint junction between *STAT3* intron 8-9 and *NBR2* intron 1-2. In the *STAT3/BRCA1* fusion transcript, the last nucleotide of *STAT3* exon 8 is fused to the first nucleotide of *BRCA1* exon 2, which carries the BRCA1 translation initiation codon (ATG, highlighted in bold red). **H)** Tumor growth of the four TNBC PDX models of the longitudinal series. The average percentage reduction in tumor growth rate for the carboplatin arm compared to the control arm is reported on each graph. Responders are identified as PDX models showing a > 5% growth rate reduction.

We further investigated the relationship between *BRCA1* methylation status and treatment exposure using four TNBC PDX models, WHIM68, WHIM69, WHIM74 and WHIM75 (i.e., the WHIM PDX longitudinal series), established from subsequent breast cancer biopsies and surgical specimens from the same patient (Figure 3D). WHIM68 was established from a pre-neoadjuvant treatment breast biopsy and it is therefore treatment-naïve, WHIM69 was derived from a research biopsy taken at day 3 of the first cycle of neoadjuvant treatment with carboplatin and docetaxel; WHIM74 was derived from a mastectomy specimen obtained after the completion of the neoadjuvant course; and WHIM75 was established from a liver metastasis. This PDX series allowed us to ask whether chemotherapeutic treatment of an individual patient would result in the progressive loss of *BRCA1* promoter methylation and in the augmentation of steady state *BRCA1* expression. We first assessed the *BRCA1* methylation status of the four serial PDXs via MSP and found that while WHIM68 was completely methylated, the two subsequent post-treatment PDX models, WHIM69 and WHIM74, showed the presence of an unmethylated signal, suggesting progressive loss of *BRCA1* methylation with exposure to therapy. (Figure 3E). *BRCA1* gene expression, as assessed by quantitative PCR, was nearly undetectable in the WHIM68 model, but progressively increased in each successive PDX model, reaching levels comparable to the BRCA1 proficient status in WHIM75 (Figure 3F). The results from the first three PDXs in the series suggest that loss of *BRCA1* methylation can occur shortly after a single exposure to chemotherapy in the patient, and that the resultant cancers progressively increase *BRCA1* expression after successive chemotherapeutic exposures. By contrast, WHIM75 appeared to be an outlier, in that it showed normal levels of *BRCA1* expression while retaining a complete methylation profile. We hypothesized that the elevated levels of *BRCA1* in WHIM75 may be caused by a form of promoter bypass and scanned its genome for somatic rearrangements engaging the *BRCA1* locus. In a detailed genomic analysis, we found a ∼800 Kb tandem duplication resulting in the fusion of the *NBR2* non-protein coding gene, which resides adjacent to the *BRCA1* promoter, with the *STAT3* gene, located several hundreds of Kb upstream of *BRCA1* (Figure S2A). We confirmed via PCR and Sanger sequencing that this rearrangement is specific to WHIM75 and is not found in any one of the other three PDX models from the same patient (Figure S2B). Sequence analysis of the breakpoint junction revealed a 4 nucleotide microhomology region, but no larger stretch of homology between the two gene fusion partners (Figure S2C). Based on the exonic structure of the genes involved in the rearrangement, we predicted that the duplication would result in the production of a fusion transcript between the *STAT3* and the *BRCA1* genes, which would place the *BRCA1* gene under the control of the *STAT3* promoter, effectively bypassing the negative regulation of the hypermethylated *BRCA1* promoter. Reverse transcription PCR and Sanger sequencing confirmed the presence of a *STAT3/BRCA1* fusion transcript, where the last nucleotide of *STAT3* exon 8 is fused to the first nucleotide of *BRCA1* exon 2, which carries the BRCA1 translation initiation codon (Figure 3G), and that its expression is specific to the WHIM75 PDX model (Figure S2D). A similar fusion has been previously reported when a TNBC *BRCA1*meth PDX was rendered resistant to cisplatin therapy in an experimental setting (Ter Brugge et al., 2016), but, to our knowledge, this is the first time that a similar mechanism of BRCA1 re-activation is identified in a clinically-derived sample. Importantly, the progressive re-expression of *BRCA1* in the WHIM PDX series, either via loss of methylation (in WHIM68 and WHIM74) or by promoter hijacking (in WHIM75) was associated with resistance to carboplatin *in vivo* (Figure 3H). The WHIM PDX series therefore showed that re-activation of BRCA1 can occur via alternative mechanisms in different *BRCA1*meth cancer tissues from a single patient, namely loss of *BRCA1* methylation at one site (i.e., the breast) and a genomic rearrangement placing the *BRCA1* gene under the control of a heterologous promoter at another site (i.e., metastasis to the liver), with both mechanisms being associated with acquired resistance to platinum-based therapy. Based on the response pattern observed across the WHIM series, we explored the possible association between the degree of *BRCA1* methylation and platinum-sensitivity across the 16 PDX models of *BRCA1*meth TNBC, each one of which has been assessed for its *in vivo* response to either cisplatin or carboplatin therapy. We found that PDXs with complete methylation of the *BRCA1* promoter had significantly higher response rates compared to those with partial methylation (82% response rate (n =11) vs. 20% response rate (n = 5), P = 0.036, **Table S5** and Figure 3C).

We then evaluated whether direct treatment with chemotherapy of a *BRCA1*meth PDX can cause loss of *BRCA1* methylation, by selecting a single completely methylated TNBC PDX established from a treatment-naïve patient cancer (TM00097) and subjecting it to treatment *in vivo* with four weekly doses of either cisplatin or vehicle control, followed by regular monitoring of tumor recurrence (Figure 4A). We found that while only one of the three tumors in the control arm showed a weak unmethylated signal, all six cisplatin recurrences showed the emergence of a strong unmethylated signal, clearly suggesting that chemotherapeutic exposure is the cause of the epigenetic shift at the *BRCA1* promoter locus (Figure 4B). Similar results were obtained with a second treatment-naïve and completely methylated PDX model (TM01079), with all three tumors in the control arm maintaining complete methylation, but five out of six cisplatin recurrences showing the emergence of the unmethylated signal (Figure 4C and D). Importantly, loss of methylation was accompanied by re-expression of the *BRCA1* gene to levels often comparable to those found in non*BRCA* and *BRCA1*mut tumors (Figure 4E). We next asked the question of whether compounds commonly used in the clinical treatment of TNBC patients, such as doxorubicin, cyclophosphamide and taxanes, can similarly cause the conversion from complete to partial *BRCA1* methylation and, consequently, restoration of *BRCA1* expression. To this end, we treated the TM00097 TNBC PDX model with a combination of doxorubicin and cyclophosphamide, administered weekly for three weeks, followed by three weekly doses of docetaxel (i.e., AC→T). The AC→T regimen, mimicking a commonly used TNBC clinical protocol, was efficient in reducing tumor volume to almost undetectable levels (Figure S3A). However, after ∼60 days of drug holiday, tumors eventually relapsed. When tested for *BRCA1* promoter methylation, all three tumor residuals from the control arm maintained the complete *BRCA1* methylation profile, while all three tumor relapses form the AC→T arm converted to the partial methylation status (Figure S3B) similar to their cisplatin treated counterparts in the previous experiments. Again, loss of methylation in the AC→T relapses was associated with a significant increase in *BRCA1* expression (Figure S3C). This states that standard non-platinum regimens commonly used in TNBC can induce the partial methylation status associated with higher *BRCA1* expression and platinum insensitivity.

**Figure 4.**
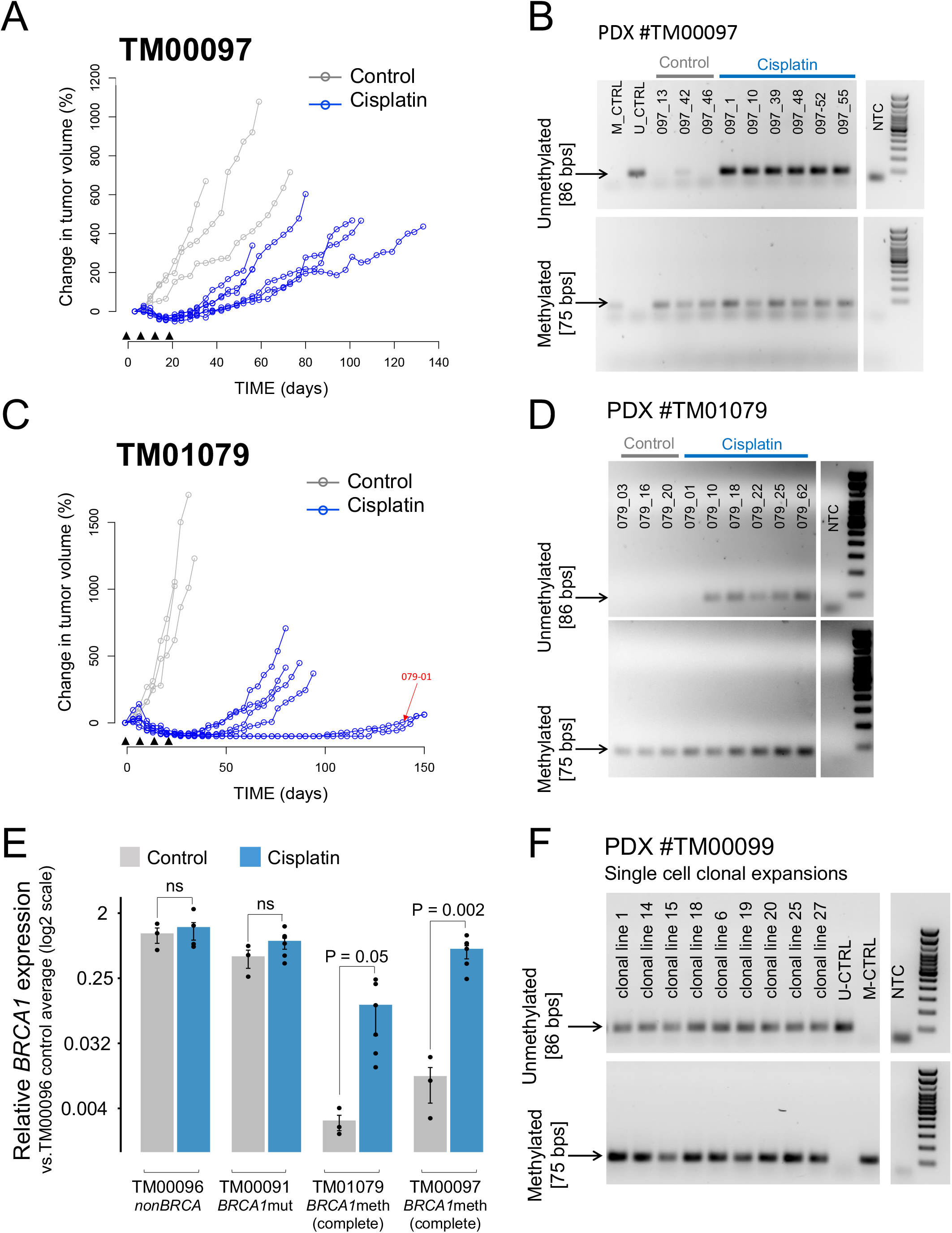
Assessment of *BRCA1* methylation and expression in TNBC PDXs treated with cisplatin *in vivo*. **A)** Tumor growth curves for TNBC PDX #TM00097. Black arrow heads indicate the timing of the four weekly doses of cisplatin. **B)** MSP results for vehicle tumors and cisplatin-treated tumor recurrences relative to PDX #TM00097. **C)** Tumor growth curves for TNBC PDX #TM01097. Black arrow heads indicate weekly doses of cisplatin. A red arrow indicates the growth curve for animal #097-01, whose recurrent tumor maintained a full methylation profile, as assessed in **(D)**. **D)** MSP results for vehicle tumors and cisplatin-treated tumor recurrences relative to PDX #TM01079. **E)** *BRCA1* gene expression (qPCR) for vehicle tumors and cisplatin- or docetaxel-treated recurrences relative to four TNBC PDX models with different *BRCA* backgrounds. All the data is normalized to the average *BRCA1* expression level of control tumors from the non*BRCA* PDX #TM00096. P values by Student’s t-test (two-tailed). **F)** MSP results for nine single cell derived clonal expansions of the primary cultures established from the *BRCA1* partially methylated TNBC PDX #TM00099.

One proposed hypothesis for the loss of *BRCA1* methylation following chemotherapeutic treatment and the emergence of resistant tumor recurrences, is the rapid expansion of a non-methylated subclone from a heterogeneous tumor cell population that contains *BRCA1*meth and non*BRCA1* clones. Patch et al. described the case of a patient (AOCS-091) initially diagnosed with a *BRCA1* methylated, platinum sensitive primary OvCa, who eventually developed a recurrent cancer that was unmethylated and chemo resistant (Patch et al., 2015). It was speculated that the recurrence may have originated from a completely independent original non*BRCA1* (i.e., *BRCA1* proficient and promoter unmethylated) and platinum resistant subclone, which was expanded during the chemotherapeutic treatment (Patch et al., 2015). Through a detailed genomic reanalysis of structural mutations in the AOCS-091 OvCa pair, we found that, despite the presence of many private rearrangements, both the primary and the recurrent cancer genomes classified as TDP type 1 (Figure S2E), indicating that the recurrent tumor originated from a subclone that, early in its evolutionary history, had experienced significant BRCA1 deficiency, most likely by promoter methylation, in order to generate BRCA1-related genomic scars before eventually losing *BRCA1* methylation at a later timepoint. To further explore whether the observed loss of *BRCA1* methylation is due to the expansion of a non-methylated tumor subclone or to active demethylation (i.e., conversion of one *BRCA1* promoter allele from the methylated to the unmethylated state), we established single cell clonal expansions from the TM0099 PDX model, a TDP type 1 TNBC with two copies of the *BRCA1* gene and partial methylation at the *BRCA1* promoter, when assessed in the bulk tumor. MSP analysis of each of the nine individual clonal lines examined showed both methylated and unmethylated amplification products (Figure 3F). This suggests that, at least in the TM0099 model, loss of methylation was the result of demethylation of one of the two *BRCA1* promoter alleles, and not of the expansion of a non-methylated subclone.

### Differential DNA methylation and gene expression between *BRCA1*meth and *BRCA1*mut cancer genomes

We have thus far established that demethylation at the *BRCA1* promoter is responsible for the recovery of *BRCA1* expression and likely leading to progressive insensitivity to platinum compounds, and to other DNA damaging agents. We asked whether methylation at the *BRCA1* promoter is associated with wider epigenetic changes that would result in the activation of critical chemoresistance genes from the onset during the establishment of the *BRCA1*meth status. To address this, we sought to identify other regions of the genome that underwent differentially methylation changes between *BRCA1*meth and *BRCA1*mut cancers across five independent TNBC and OvCa methylation array datasets (**Table S6** and Figure S4A). We first performed a probe-wise differential methylation analysis to identify individual CpG sites that are differentially methylated between the two *BRCA1* states. All the differentially methylated CpGs identified across the five independently analyzed datasets, mapped to a ∼80 Kb region centered around the *BRCA1* promoter, with a large majority of the significant CpGs (102/111, 91.9%) clustered exactly at the *BRCA1* promoter region (Figure S4B). We further confirmed that significant changes in DNA methylation between BRCA*1*meth and *BRCA1*mut cancers are restricted precisely to the *BRCA1* promoter, by identifying differentially methylated genomic regions using the *bumphunter* function of the *minfi* R package: the only region with a significant change in methylation patterns between the two groups corresponded to the *BRCA1* minimal promoter (chr17:41,277,059-41,278,506, hg19, Figures S4C). This indicates that hypermethylation at the *BRCA1* promoter is an extremely localized event and does not reflect more widespread epigenetic changes. Similarly, when we performed an analysis of differential gene expression between *BRCA1*meth and *BRCA1*mut cancers across the same datasets, we found *BRCA1* and *NBR2* (which is under the control of the same methylation-sensitive bidirectional promoter as *BRCA1*) as the only two significantly differentially expressed genes (Figure S4D). These results indicate that epigenetic silencing of the *BRCA1* gene is an isolated event in the cancer genome whose unique consequence is the loss of BRCA1 activity reflected by the emergence of genomic scars indistinguishable from those driven by pathogenic mutations in the *BRCA1* gene. This is in agreement with a recent study by Glodzik *et al*. that focuses only on TNBC (Glodzik et al., 2020), but our results extend this observation to OvCa.

Taken together, these data suggest that BRCA1 deficiency due to promoter methylation can be overridden by demethylation of one allele associated with restoration of *BRCA1* mRNA expression, which can occur after only a short course of platinum chemotherapy both in TNBC and in OvCa. Finally, once the partially methylated state is established, the resultant cancer exhibits relative resistance to platinum drugs akin to non*BRCA* cancers, which appears to be mediated by *BRCA1* re-expression.

### *BRCA* gene mutations, and not *BRCA1* methylation, are also strongly predictive of platinum-based therapy response in independent OvCa cohorts

Previously we determined that BRCA1 deficiency exhibits the identical effect on inducing the TDP in both TNBC and OvCa (Menghi et al., 2018). If the primary genomic biology of TDP formation is the driver for this chemotherapeutic response, then it should hold across different disease types with the same genomic characteristics. We sought to validate the hypothesis that *BRCA* mutational status and not TDP status is predictive of patient response to platinum-based chemotherapy in OvCa cohorts.

In the first instance, we looked to publicly available data for ovarian cancer cohorts with complete genome sequence information and response assessment. To this end, the only dataset with detailed response information along with whole genome sequencing and a high number of cases (n = 80 primary cancers) was from the Australian Ovarian Cancer Study (AOCS (Patch et al., 2015), Figure 5A, **Tables S1-2**). Despite the genomic similarities between TDP OvCa and TNBC cancers, the presenting state of the cancers, especially in terms of tumor burden, and the assessment of response are very different. At presentation, OvCa has significantly higher tumor burden than TNBC, and the determination of response in OvCa (as compared to TNBC) is not primarily by quantitative assessment of tumor shrinkage (such as assessment of pCR) but by duration of response and overall survival. The AOCS cohort included 32 TDP type 1 cancers, 30 of which exhibit some form of BRCA1 abrogation (Figure 5B). When OvCa patients were stratified based on their TDP status there was no significant association with survival (Figure S5A). However, patients with *BRCA*mut cancers showed a significant improved overall survival compared to those with non*BRCA* cancers (Hazard Ratio (HR) = 0.52, P = 0.025; Figure 5C), while, again, *BRCA1* promoter methylation did not provide any significant benefit. None of the other available clinical variables were linked to better survival in a univariate analysis (**Table S3**), but age became significant in the multivariate model correcting for *BRCA* status, a trend driven by the strong association between *BRCA1* methylation status and younger patient age (Figure S5B).

**Figure 5.**
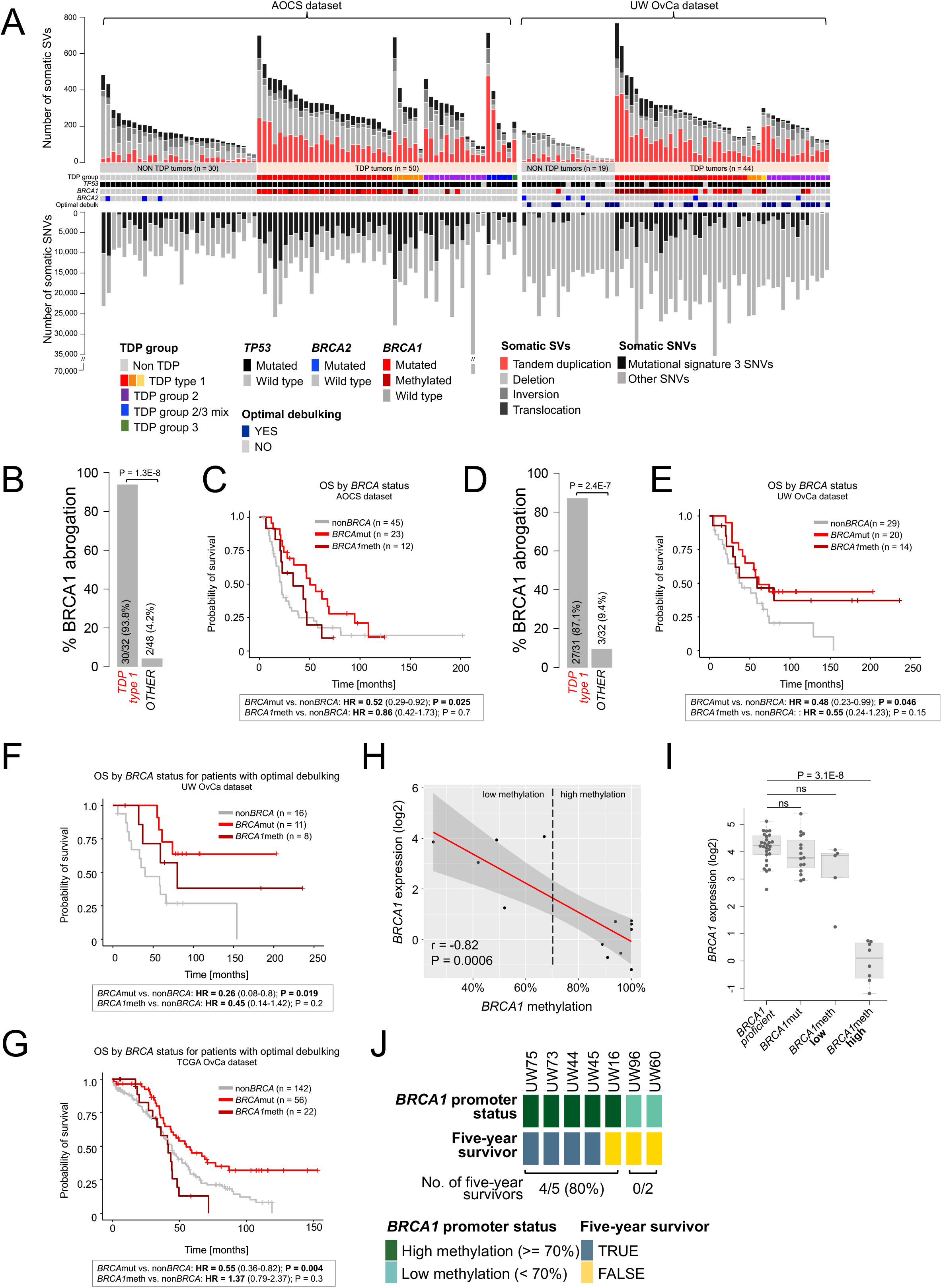
Genomic features and therapeutic response across independent cohorts of OvCa patients. **A)** Overview of the genomic and clinical features relative to the AOCS and the UW OvCa cohorts. Samples are sorted based on their TDP group assignment. **B)** AOCS cohort: percentage of tumors with BRCA1 abrogation relative to TDP status. P by logistic regression. **C)** AOCS cohort: overall survival (OS) as a function of *BRCA* status (HR, hazard ratio; CI, confidence interval; P value by Cox proportional hazards regression model). **D-E)** Same as B-C, relative to the UW OvCa cohort. **F)** UW OvCa cohort: overall survival as a function of *BRCA* status, including only optimally debulked patients. **G)** TCGA OvCa cohort: overall survival as a function of *BRCA* status, including only optimally debulked patients. **H)** Scatter plot of *BRCA1* gene expression (RNAseq log 2 values) and *BRCA1* promoter methylation (based on MS-ddPCR estimates corrected for the proportion of neoplastic cellularity) for the subset of 13 *BRCA1* methylated UW OvCas with available expression data. A smooth local regression line with 95% confidence interval was drawn using the *geom_smooth* function in R (method = loess). The Pearson correlation coefficient (r) and its corresponding p-value (P) are shown. **I)** *BRCA1* gene expression across the UW OvCa cohort as a function of *BRCA1* status. P value by Student’s t-test (two-tailed). **J)** *BRCA1* methylation status (via MS-ddPCR) and five-year survivorship for the subset of optimally debulked patients with *BRCA1*meth cancers in the UW OvCa cohort. Patients with fewer than 5 years of follow up are not included.

Given the positive signal from the publicly available data, we pursued more in-depth analysis in an independent OvCa dataset. To this end, we analyzed a new cohort of 63 primary OvCa patients with detailed response data to therapy with a combination of carboplatin and paclitaxel, from the University of Washington Medical Center (UW OvCa cohort, Figure 5A, **Tables S1-2**). In this cohort, BRCA1 deficiency was again significantly associated with the TDP type 1 configuration (27/31 (87.1%) vs. 3/32 (9.4%) for non TDP type 1 cancers, P = 2.4E-7, Figure 5D), with similar proportions of *BRCA1*mut and *BRCA1*methy OvCas defining the BRCA1 deficient group (**Table S2**). When we examined the correlation of overall survival with *BRCA* status in the UW OvCa cohort, again, we found that patients with *BRCA*mut OvCas, but not those with *BRCA1*meth OvCas, had significantly better overall survival when compared to patients with non*BRCA* OvCas (HR = 0.48, P = 0.046, Figure 5E). Since optimal debulking also appeared to be associated with survival in a univariate analysis in this dataset (Figure S5C and **Table S3**), we separated patients who were optimally (defined as maximum residual tumor diameter < 1 cm) or sub optimally debulked. *BRCA* mutational status highly correlated with better overall survival exclusively in the subgroup of patients who were optimally debulked (HR = 0.26, P = 0.019, Figures 5F and S5D, and **Table S3**). Importantly, in this patient subgroup, *BRCA1* methylation showed an intermediate effect that did not reach statistical significance (Figure 5F), while TDP status did not associate with improved survival, regardless of optimal debulking (Figures S5C-D). Taken together, the analysis of the two OvCa clinical cohorts and of the COH TNBC cohort show that *BRCA* mutation status but not *BRCA1* methylation, nor TDP status is consistently predictive of platinum response and that these associations are consistent across the two cancer types examined.

Adequacy of surgical cytoreduction is a known powerful prognostic factor in OvCa outcomes. Since the AOCS dataset did not have debulking information, we sought another large data set upon which we could validate the interaction of *BRCA* status and overall survival, while controlling for adequacy of surgical cytoreduction. The TCGA ovarian cancer cohort comprises 314 primary high grade serous ovarian carcinomas from patients that went on to receive carboplatin-based chemotherapy and with known *BRCA* status and, in the majority of cases, debulking status (Cancer Genome Atlas Research, 2011). Survival analysis of the full dataset confirmed a significant association between *BRCA* mutation and better overall survival (HR = 0.053, P = 0.0003; Figure S5E), which was maintained after correcting for several outcome-associated clinical variables, including optimal debulking, race and age (**Table S3**). As expected, debulking status was the clinical variable most significantly associated with overall survival in the full dataset (Figure S5E). When we split the TCGA cohort based on debulking status, we found that patients with *BRCA*mut OvCas had better overall survival whether they were optimally or sub optimally debulked (*BRCA*mut vs. non*BRCA*; HR = 0.55, P = 0.004 for optimally debulked patients; HR = 0.47, P = 0.051 for sub optimally debulked patients; Figures 5G and S5E, **Table S3**). By contrast, *BRCA1* methylation status did not provide any benefit in terms of overall survival, even in the subgroup of optimally debulked patients. Thus, like in TNBC, *BRCA* deficiency via disruptive mutations, but not *BRCA1* promoter methylation, was significantly associated with improved response to platinum-based therapy in OvCa.

In the UW OvCa cohort, we had the opportunity to examine the consequences of the degree of *BRCA1* promoter methylation on gene expression directly in a patient cohort, as a validation of the results obtained in the PDX TNBC cohort. To this end, we employed methylation-specific droplet digital PCR (MS-ddPCR) to reassess the *BRCA1* methylation status of the cancer genomes in the UW OvCa cohort in a quantitative manner (Kondrashova et al., 2018). After correcting for the proportion of neoplastic cellularity, the degree of *BRCA1* methylation showed a strong and significant negative correlation with *BRCA1* expression levels (r = −0.82, P = 0.0006, Figure 5H). This analysis also showed that *BRCA1*meth OvCas can be separated into two groups by setting a quantitative methylation threshold of 70% to identify cancers with low-methylation (n = 5) and *BRCA1* expression levels akin to those found in non*BRCA1* cancers (median fold change = 0.77, not significant), and cancers with high-methylation (n = 8) with *BRCA1* expression levels reduced to less than 10% of those observed in non*BRCA1* cancers, (median fold change = 0.06, P = 3.1E-8, Figure 5I). Interestingly, this methylation threshold is identical to one previously proposed for OvCa in a different but larger cohort (Kondrashova et al., 2018). The modest number of patients in the high methylation group did not permit a proper statistical assessment of the clinical benefit associated with this profile in terms of prolonged overall survival. However, when we stratified patients based on their *BRCA1* methylation profile and adequate surgical cytoreduction, we found that 4/5 patients with high methylation survived five years or longer, but neither of the two patients with low methylation did (Figure 5J). While the numbers are too small to assess statistical significance, this trend, specific for the subset of patients who were optimally debulked, is consistent with our observations in the PDX TNBC cohort.

### In patients with non*BRCA* TNBC and OvCa, optimal response to the platinum/taxane chemotherapeutic combination is associated with an enhanced immune signature

We next sought to identify features associated with clinical response for patients with non*BRCA* cancers, starting with the COH TNBC cohort. Because we found no gene alterations, other than those affecting the *BRCA* genes, that were predictive for chemo-responsiveness in this cohort, we looked for transcriptional signals that associated with sensitivity to chemotherapy. The TNBCtype classification described by Chen et al. segregates TNBCs into six transcriptional subsets, including a specific immunomodulatory subtype characteristic of TNBCs with a high level of immune cell infiltrates (Chen et al., 2012). Interestingly, we saw a significant enrichment for the immunomodulatory subtype in cancers from patients who achieved pCR (7/15 (46.6%) vs. 1/18 (5.5%), OR = 13.6, P = 0.01, Figure 6A), but not for other expression-based subtypes. We then looked for specific expression profiles associated with better response across the entire TNBC cohort and found that the genes significantly over-expressed in TNBCs from patients with pCR were highly enriched for immune response-related pathways, such as adaptive immunity and interferon signaling (Figure S6A), with a highly significant proportion of over-expressed genes annotated as immune genes (p = 6.0E-76, Figure S6B). When we sub-grouped TNBCs based on their *BRCA* status, the significant association between increased expression of immune genes and a higher pCR rate was only found in the non*BRCA* TNBC subset, but not in the *BRCA1*meth subset (Figure 6B). This suggests that the detected immune signal may be specifically linked to better response on a *BRCA*-proficient background. Because of the small sample size of the non-pCR subset of patients with *BRCA*mut TNBCs (n = 1), we could not perform an equivalent analysis of differential gene expression in the *BRCA*mut subgroup.

**Figure 6.**
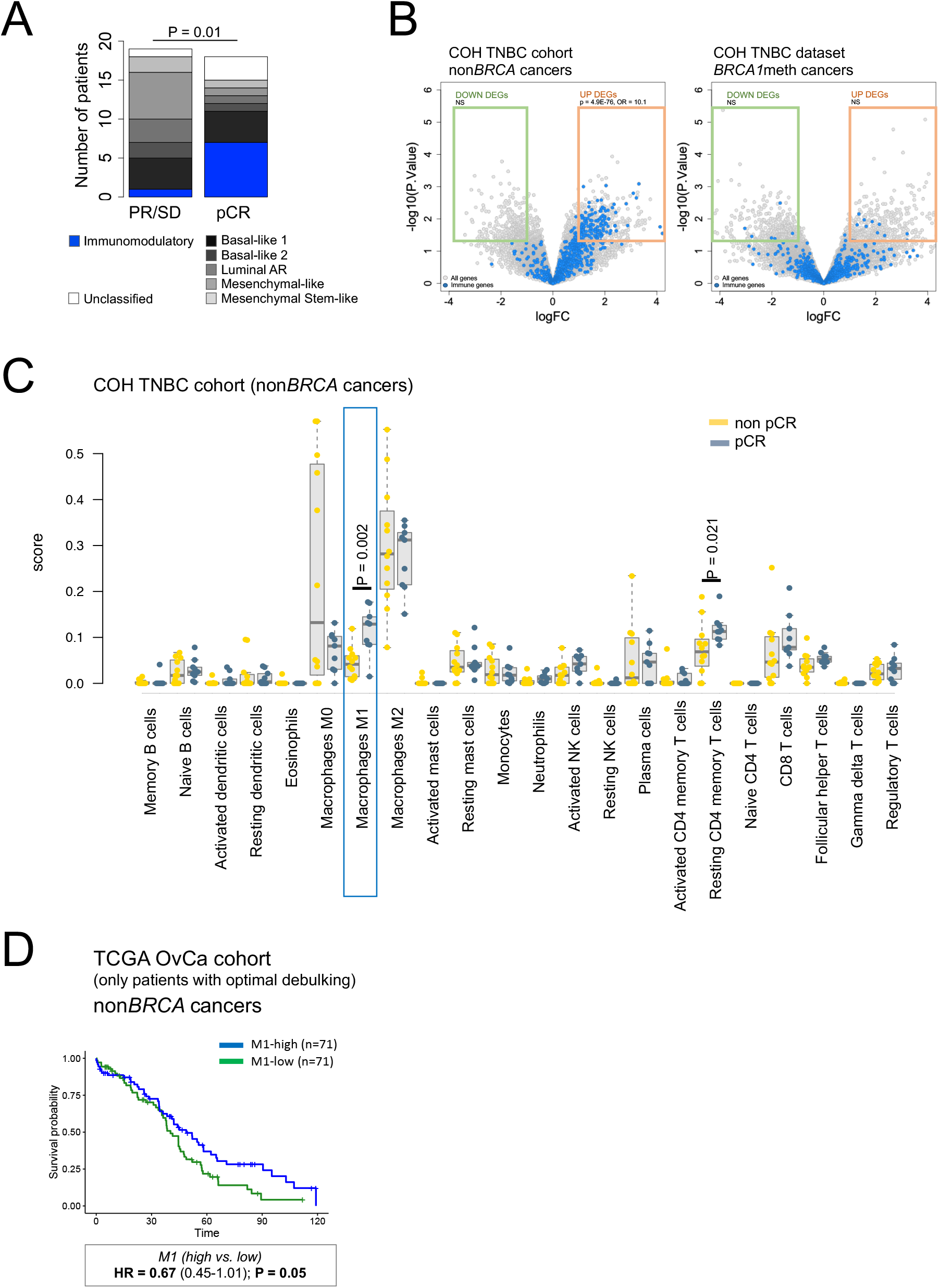
Immune expression signals and therapeutic response across *BRCA1/2* wild type tumors. **A)** TNBCtype classification of the COH TNBCs based on the six transcriptional subtypes described by Chen et al (Chen et al., 2012), showing a significant enrichment of immunomodulatory subtype among TNBC patients who achieved pCR. P value by Fisher’s exact test. Unclassified tumors are depicted in white and were not included in the statistical analysis. **B)** Volcano plots of differential gene expression between COH TNBC patient who achieved pCR and those who did not. Significantly differentially expressed genes (p-value < 0.05 and absolute log2 fold change > 1), are boxed in green (DOWN) and orange (UP). Immune genes are shown in blue. Enrichment of immune genes within the down- and up-regulated differentially expressed genes was computed by Fisher’s exact test. **C)** Comparison of cancer CIBERSORT scores between patients who achieved and did not achieve pCR, in the COH TNBC cohort, using the Mann-Whitney test. Only significant P values are reported. M1 macrophage scores highlighted by a blue box. **D)** Overall survival analysis of optimally debulked TCGA OvCa patients with non*BRCA* cancers. Patients are stratified into M1-high and M1-low, based on their CIBERSORT-derived M1 macrophage signature score (split on the median level for the group). HR, hazard ratio; CI, confidence interval; P value by Cox proportional hazards regression model.

We then asked if the observed immune signal could be parsed out into different types of infiltrating immune cells. To this end, we applied the CIBERSORT computational approach and assessed the relative abundance of 22 distinct immune cell types for each TNBC in the COH TNBC cohort with RNAseq expression profiling. The most significant association from the comparison of the CIBERSORT-generated scores between pCR vs. non-pCR subgroups was an increase in M1 macrophage scores (Figure 6C). Similar to the genetic analysis of differential gene expression described above, this association was not found when analyzing the subset of *BRCA1*meth TNBCs (Figure S7A). We further investigated the association between higher M1 macrophage scores and better chemotherapeutic response, through an analysis of overall survival across the three OvCa cohorts. In both the AOCS and the UW OvCa cohorts, patients with non*BRCA* OvCas and higher M1 macrophage scores (i.e., M1-high, split on the median value) showed a trend for longer overall survival, which, however, did not reach statistical significance (Figures S7B). We hypothesized that lack of significance in these datasets may be due to the potential confounding effects of optimal debulking, and relatively small sample size. To assess this, we examined the TCGA OvCa dataset. When the analysis was limited to the subset of optimally debulked patients, a significant association between M1-high scores and longer overall survival was observed only in patients with non*BRCA* OvCas (Figure 6D), but not in those with either *BRCA*mut or *BRCA1*meth OvCas (Figures S7C-D). As the association of specific immune expression phenotypes with better survival in OvCa has been previously investigated with contrasting outcomes (Cancer Genome Atlas Research, 2011; Liu et al., 2020; Thorsson et al., 2018; Verhaak et al., 2013), our analyses suggest that the beneficial effect of an augmented immune signature may be limited to non*BRCA* cancers, and potentially driven by M1 macrophage effects.

### A proposed decision tree for predicting response to the platinum/taxane chemotherapeutic combination in TNBC and OvCa

Our results strongly suggest that by considering *BRCA* status, as well as the expression of an immune signature in non*BRCA* cancers, we might be able to better predict chemo-responsiveness than by using *BRCA* mutational status alone or by assessing the HRD status in both TNBC and OvCa. To test this, we developed a combined predictor that could better correlate with outcomes to platinum-based chemotherapy based on the presence of *BRCA* pathogenic mutations and, in non*BRCA* cancers, on the strength of the M1 macrophage transcriptional signal as indicated in the schematic in Figure 7A. We compared the predictive performance of our combined response criteria with that of *BRCA* status alone and of a surrogate measure of HRD, i.e., HRD proxy, that combines TDP type 1 and *BRCA2* mutant tumors into an HRD proxy high category. When applied to the COH TNBC cohort, 84.2% (16/19) of the patients predicted to have optimal therapeutic outcomes based on our combined response criteria did achieve pCR. On the other end, only 9.1% (1/11) of TNBC patients predicted to be poor responders achieved pCR. When considered altogether, the accuracy of our algorithm in predicting pCR was 81% (Figure 7B). By contrasts, the predictive accuracy of *BRCA* status alone and of the surrogate for HRD were 69% and 56%, respectively (Figure 7B). We then compared the three modes of patient stratification across the three OvCa cohorts examined in this study, using Cox proportional hazard ratio test statistics to examine overall survival trends. In each of the three cohorts, the response outcomes predicted by our proposed combined response criteria outperformed the predictive power of *BRCA* status alone and of HRD/HRD proxy metrics, as indicated by improved P values and lower hazard ratios (Figure 7C). More importantly, the number of cases predicted to be good responders increased significantly. For example, for the TCGA OvCa cohort, this number increases by over 2-fold from 48 to 110 individuals predicted to have good therapeutic outcomes based on either their *BRCA* status or the combined response criteria, respectively (Figure 7C). These findings further support our hypothesis that BRCA deficiency status (both *BRCA1* and *BRCA2*) computed via genome-based measures of HRD is not a good predictor of chemotherapeutic response *per se*, and that additional patient stratification focused on *BRCA* mutation/methylation status and immune transcriptional profiles, and the M1 macrophage signature in particular, could provide improved clinical prediction in TNBC and OvCa if prospectively validated.

**Figure 7.**
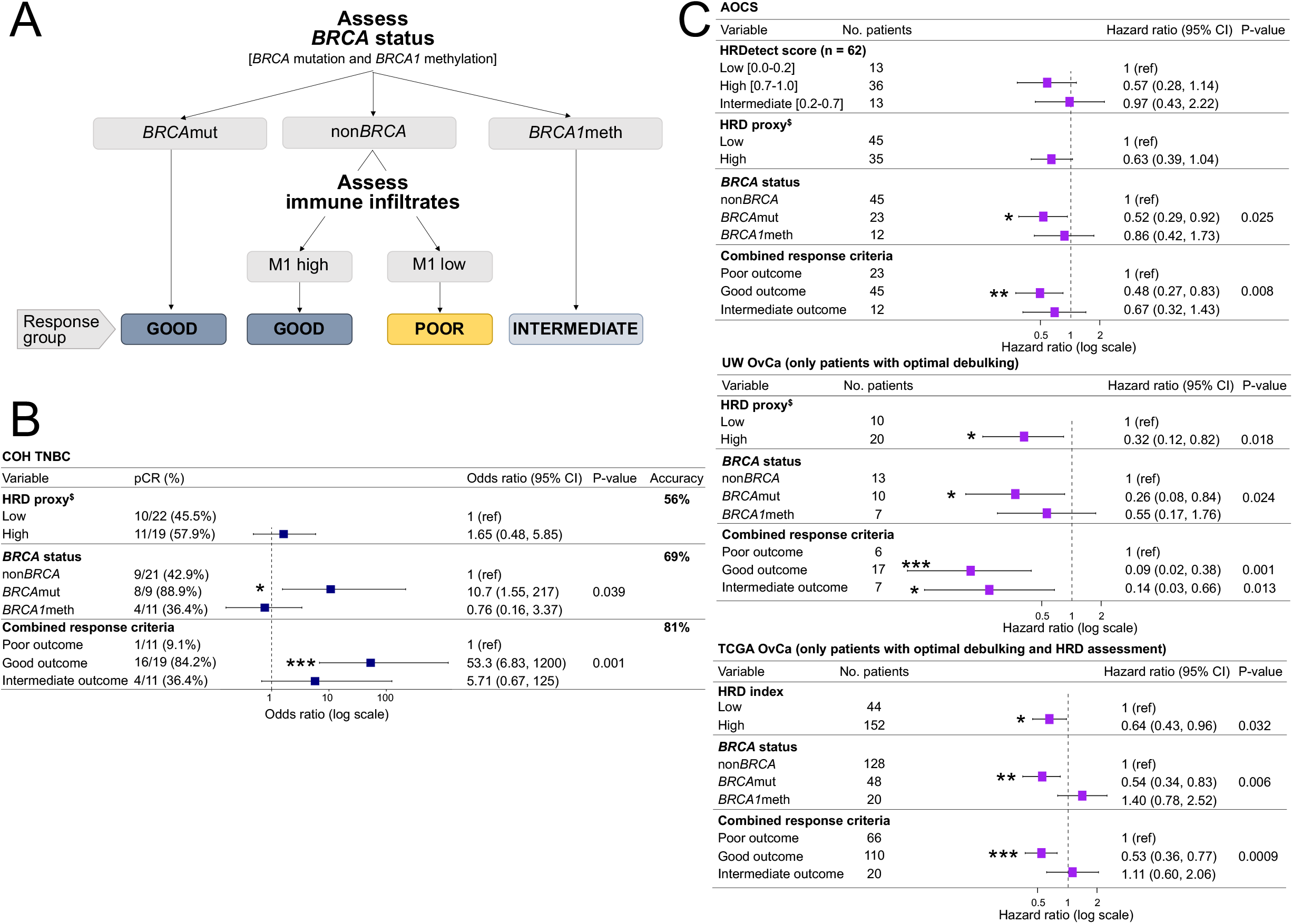
A decision tree for predicting response of TNBC and OvCa patients to platinum-taxane combination chemotherapy. **A)** Schematic of the proposed decision tree for optimal prediction of TNCB and OvCa patient response to platinum-taxane combination chemotherapy. M1 refers to the macrophage M1 signal computed using CIBERSORT, with high and low defined based on the median score value for the non*BRCA* subset of cancers**. B)** Odd ratios of pCR for the COH TNBC cohort, with patients stratified based on either *BRCA* status alone, HRD low vs. high, or the combined response criteria (computed as described in (**A**)) Odds ratio with 95% confidence intervals and p-values were computed using logistic regression. **C)** Hazard ratios for overall survival of OvCa patients for the UW OvCa, AOCS and TCGA OvCa cohorts, stratified as described in (**B**)). For the AOCS cohort, patient stratification based on HRDetect scores as reported by Davies et al. **(Davies et al., 2017)** is also reported for comparison. Hazard ratio with 95% confidence intervals and p-values were computed using the COX proportional hazard test. ^$^HRD proxy refers to an estimate of HRD which combines TDP type 1 cancers and *BRCA2* mutated cancers into the HRD proxy high category. * P < 0.05; ** P < 0.01; *** P < 0.001.

## DISCUSSION

A major focus of this study was to explore the root causes for the inconsistencies in the clinical data when defining chemotherapeutic response in TNBC and OvCa. Earlier, it was thought that all forms of BRCA deficiency were sufficient to render a cancer sensitive to DNA damaging agents such as platinum compounds. This led to the use of measures of BRCA deficiency through the assessment of BRCA-related genomic scars (i.e., HRD scores) to project chemotherapeutic responsiveness especially in OvCa (Abkevich et al., 2012; Gorodnova et al., 2015; Popova et al., 2012; Stronach et al., 2018; Telli et al., 2016; Watkins et al., 2014). Subsequent clinical studies suggest that although measures of HRD are good indicators of BRCA1 and BRCA2 deficiency, they are not reliable tools to assess chemotherapeutic responsiveness (Isakoff et al., 2015; Sharma et al., 2018; Staaf et al., 2019; Stronach et al., 2018; Telli et al., 2016; Tutt et al., 2018; Watkins et al., 2014). A comprehensive meta-analysis of 34 studies encompassing 7,986 treated OvCa patients showed that *BRCA*mut cancers were significantly associated with better survival compared to nonBRCA cancers, but that *BRCA1*meth cancers had no such association (Sun et al., 2014). In the two-arm randomized TNT clinical trial, Tutt et al. reported that, contrary to *BRCA1* germline mutations, neither *BRCA1* methylation nor Myriad HRD assessment were associated with better response to carboplatin compared to docetaxel in metastatic TNBC (Tutt et al., 2018). Our work herein supports this growing number of reports in the clinical literature showing that *BRCA1*meth cancers have significantly lower objective response to platinum compounds than *BRCA1*mut cancers in all clinical TNBC and OvCa cohorts examined and in the PDX model dataset. This happens despite the fact that *BRCA1* promoter methylation is associated with a significant and dramatic reduction in *BRCA1* RNA levels and with patterns of TDP type 1 formation identical to those found in *BRCA1*mut cancer genomes. One hypothesis for the lack of a survival advantage in patients with *BRCA1*meth OvCas is the rapid selection of a non-methylated subclone following platinum-treatment of a heterogenous tumor cell population (Patch et al., 2015). However, the emergence of a completely unmethylated clone as the only mechanism is not a consistent observation (Swisher et al., 2017). Furthermore, using mouse models of mammary tumors driven by the loss of *Trp53* and *Brca1*, we previously showed that both copies of the *Brca1* gene must be lost for the TDP type 1 configuration to emerge (Menghi et al., 2018). We therefore postulate that TDP genomes must have experienced a full loss of BRCA1 activity and that a hidden clone with heterozygous *BRCA1* methylation would not be sufficient to induce the TDP type 1. Thus, the emergence of a clone from a TDP tumor that was partially methylated at the *BRCA1* promoter is highly improbable.

Our results strongly suggest, however, that *BRCA1* promoter methylation is a highly dynamic epigenetic state which is rapidly and consistently modified in response to even short exposure to chemotherapy. We show with a variety of PDX models of *BRCA1*meth TNBC that, indeed, following only one cycle of platinum treatment, cancer genomes with complete *BRCA1* methylation convert primarily to a partial *BRCA1*meth state where both methylated and unmethylated promoter alleles coexist leading to an increased steady state expression of *BRCA1*. Our PDX longitudinal series derived from successive specimens collected before, during and after neoadjuvant chemotherapy in a single patient, confirms that exposure of a completely methylated TNBC to just a single dose of carboplatin chemotherapy results in conversion to partial methylation, enhanced *BRCA1* expression and resistance to carboplatin when assessed in the PDX model system. The speed of conversion was confirmed when fully *BRCA1* methylated PDX models were treated *in vivo* with a single course of a platin. That this partial methylated state leads to resistance is corroborated by the fact that the post-treatment derived PDX models with partial methylation showed consistent insensitivity to platinum. Our clonal analysis of primary cell cultures from a PDX model with the co-existence of a methylated and unmethylated *BRCA1* promoter showed that this configuration is due to heterozygous methylation at the *BRCA1* promoter in all individual cell clones rather than because of clonal heterogeneity. When taken together, our data suggest that *BRCA1* methylation is a functionally “plastic state” that can be rapidly lost upon chemotherapy exposure resulting in the associated restoration of *BRCA1* expression. When compounded over a series of cycles, this reversion to a *BRCA1* wild type functionality is likely to be the cause of the relative insensitivity of *BRCA1*meth cancers observed both in the experimental and clinical settings.

One of the challenges in extrapolating chemosensitivity in clinical studies is that the therapeutic regimens are almost always combinations of chemotherapeutic compounds, so the assignment of a biomarker with a specific response to a single chemotherapeutic entity cannot be formally discerned. Indeed, all the clinical cohorts analyzed herein used combination chemotherapy, although we specifically chose cohorts that exclusively used a platinum based therapeutic agent in combination with a taxane compound. To determine whether the *BRCA*-linked sensitivity is to the platinum or the taxane components of the combo regimens, we made use of ongoing single agent studies on 33 PDX TNBC models and showed significantly higher differential *in vivo* response rates to carboplatin, but not to docetaxel, for *BRCA*mut PDXs when compared to non*BRCA* PDXs. To further reduce any confounding effects of the genetic backgrounds across a range of tumors, we definitively found that the parental SUM149 cell line, which carries a pathogenetic *BRCA1* mutation, was significantly more sensitive to cisplatin and to other DNA damaging agents such as doxorubicin, and mafosfamide, compared to its BRCA1-restored SUM149.B1.S* derivative line, but with no differential sensitivity to docetaxel. We conclude therefore that the enhanced sensitivity of *BRCA*mut cancers is associated primarily with DNA damaging agents such as platinum salts, and not with a common therapeutic partner, a taxane, similar to what observed in the TNT clinical trial (Tutt et al., 2018).

One of our goals was to better resolve some conundrums posed in the clinical literature. When *BRCA1* status has been examined relative to chemosensitivity in TNBC, there have been conflicting reports of the benefit of adding platinum to a neoadjuvant anthracycline and taxane-based regimen (i.e., ACT) in patients with TNBC (Hahnen et al., 2017; Loibl et al., 2018; Sikov et al., 2015). In particular, the INFORM randomized clinical trial showed equivalent pCR rates in *BRCA1* carriers with HER2-negative breast cancers when given either neoadjuvant cisplatin or a combination of doxorubicin and cyclophosphamide (AC) (Tung et al., 2020). Our *in vitro* results suggest that depending on the doses used, the AC combination can have the equivalent BRCA1-dependent effect as a platinum compound and would explain the current clinical observations.

While we were primarily focused on the BRCA1 status question, we also found evidence that the impact of molecular heterogeneity of the *BRCA* status on therapeutic sensitivity extends to the previously observed association of immune signatures and therapeutic outcome. Several studies have reported that high immune activity measured by a range of metrics is associated with better chemotherapeutic response in both TNBC and OvCa (Kwon, 2019; Thorsson et al., 2018; Verhaak et al., 2013). However, again, these associations have never been sufficiently consistent to make this a useful predictive marker. Our analysis revealed that the salutary effects of a high immune score rested primarily in the subgroup with non*BRCA* cancers in both TNBC and OvCa. Using computational deconvolution (CIBERSORT) of the RNA seq data, we found evidence that the most consistent immunological change was the elevation of the M1 macrophage expression cassette in good responders. The M1 phenotype typically reflects classically activated tumor associated macrophages that can be induced by interferon γ and tumor necrosis factor α. M1 macrophages are believed to exert a cytotoxic effect on tumor cells, through the release of oxygen species, nitrogen intermediates and inflammatory cytokines (Zheng et al., 2017). The observed association between a stronger M1 macrophage signal and better clinical outcome is in alignment with recent evidence showing that M1 macrophage polarization in the tumor microenvironment associates with the best prognostic outcome for OvCa patients undergoing conventional surgery and chemotherapy (Maccio et al., 2020). It is not clear why this immunological effect on therapeutic outcome is particularly significant in non*BRCA* cancers since this immunological signature appears also in BRCA1 deficient cancers. One possibility is that the benefits of BRCA deficiency in chemosensitivity outweighs any benefit accrued through an immunological mechanism. Regardless of the mechanism, using this stratification of *BRCA* status and immune configuration, we constructed a putative predictive model for response to platinum in TNBC and OvCa that outperforms the use of HRD metrics, or of simply ascertaining *BRCA*mut status. At a more conceptual level, our study underscores the importance of genomic stratification in order to deconvolute phenotypic heterogeneity within clinical cohorts so that an optimal predictive diagnostic can be derived. If validated, this approach potentially identifies in advance the group of women who will have an optimal response to platinum-based chemotherapy in both TNBC and OvCa. Given that the combination of checkpoint inhibitors and chemotherapy may become the recommended neoadjuvant regimen in TNBC despite a significant increase in adverse drug events, the ability to deescalate a toxic combination in a significant subset of TNBC patients would have a meaningful impact on cancer survivors.

## AUTHOR CONTRIBUTION

Conceptualization, F.M. and E.T.L.; Methodology, F.M., R.S., P.K., I.V.R. and E.T.L.; Software and Formal Analysis: F.M., A.S.Z. and H.C.; Investigation, F.M., R.S., P.K., M.R.R., S.E.Y. and I.V.R.; Writing – Original Draft, F.M. and E.T.L.; Writing – Review & Editing, F.M., E.T.L., M.T.L., L.D., Y.Y., G.S., K.B. and E.M.S.; Funding Acquisition, F.M. and E.T.L.; Resources, Y.Y., G.S., M.T.L., L.D., K.B., and E.M.S.; Supervision, E.T.L.

## ACKNOWLEDGMENTS

RNAseq library preparation and sequencing were performed by JAX Cancer Center Shared Resources (Genomic Technology and Computational Sciences) at The Jackson Laboratory for Genomic Medicine, CT 06030, USA. WGS library preparation, sequencing and analysis were performed at the New York genome Center, New York, NY. This work was supported by NCI grant P30CA034196 (to E.T.L.), DoD CDMRP grant W81XWH-17-1-0005 (to E.T.L.), the Andrea Branch and David Elliman Cancer Study Fund (to E.T.L.), a generous gift from the Scott R. MacKenzie Foundation (to E.T.L.), an NIH/NCI PDX Trial Center for Breast Cancer Therapy grant (1U54CA22407 to M.T.L.), NIH-NCI grant U24CA22611 and NIH/NCI developmental projects grant P50CA186784 (to M.T.L.), an NIH/NCI Comprehensive Cancer Center support grant (PI: C. Kent Osborne), a generous gift from the Korell family for the study of triple-negative breast cancer (to M.T.L), funding from the Department of Defense Ovarian Cancer Research Program (OC160274, to E.M.S.) and from the National Institute of Health (R01CA237600, to E.M.S.). The COH TNBC clinical trial was funded by Celgene and supported by the City of Hope Comprehensive Cancer Center Pathology Research Services Core and Biostatistics and Mathematical Modeling Core (National Cancer Institute of the National Institutes of Health under award P30CA033572, to Y.Y. and G.S.).

## STAR METHODS

### CONTACT FOR REAGENTAND RESOURCE SHARING

Further information and requests for resources and software should be directed to and will be fulfilled by the Lead Contact, Edison T. Liu.

## METHOD DETAILS

### Tumor cohorts and WGS

Three new human tumor WGS datasets were generated as part of this study, as listed in **Table S2**. All investigations involving human specimens were performed after approval by the Institutional Review Board at each institution, and all subjects provided voluntary written informed consent.

- *City of Hope National Medical Center (COH TNBC) cohort*. Female patients with pathologically confirmed diagnosis of locally advanced and inflammatory TNBC were enrolled in a phase II trial testing the safety and efficacy of carboplatin and NAB-paclitaxel in the neoadjuvant setting (NCT01525966, (Yuan et al., 2021)) at the City of Hope Comprehensive Cancer Center. Specimens from pre-treatment tumor biopsies were snap-frozen in RNA-later solution. DNA and RNA were isolated from the same tissue fragment using the QIAGEN AllPrep DNA/RNA Mini Kit.
- *PDX cohort*. This dataset comprises 33 PDX TNBC models previously established by Prof. Micheal T. Lewis at the Baylor College of Medicine, with known response to single agent carboplatin and docetaxel, assessed as part of a PDX preclinical study (https://pdxportal.research.bcm.edu/, to be described in detail elsewhere). DNA was isolated from snap-frozen tumor tissue fragments using a QIAGEN AllPrep DNA/RNA Mini Kit.
- *University of Washington (UW OvCa) cohort*. A total of 63 serous ovarian carcinomas were selected from the Gynecologic Oncology Tissue Bank established by Prof. Elizabeth M. Swisher at the University of Washington, Seattle, WA, to represent diverse outcomes of patient survival as well as the three different *BRCA1/2* states examined in this study, but independently of any other genetic or clinical features. Patients were treated with a combination of carboplatin and paclitaxel. DNA and RNA were isolated as previously described (Bernards et al., 2018).

WGS libraries were generated using either a KAPA Hyper Prep Kit (COH TNBC cohort) or a TruSeq DNA PCR -Free kit (UW OvCa and PDX cohorts), according to manufacturer guidelines and 150 bp paired-end sequence reads were generated using either the Illumina HiSeq X Ten system (COH TNBC cohort) or a NovaSeq 6000 system (UW OvCa and PDX cohorts). Raw sequencing data were aligned to the human GRCh37 reference genome using the Burrows-Wheeler Aligner (BWA) (Li and Durbin, 2009). In the case of the PDX cohort, potential mouse contaminant reads were removed by aligning the data to a combined reference genome of mouse (GRCm38/mm10) and human (GRCh37). Structural variant calls were generated using three different tools (Crest (Wang et al., 2011), Delly (Rausch et al., 2012), and BreakDancer (Chen et al., 2009)), and high confidence events were selected when called by at least two tools and by requiring split-read support. In the absence of matched normal DNA samples to be used as controls, germline variants were identified as those that appear in the Database of Genomic Variants (DGV, http://dgv.tcag.ca/) and/or the 1,000 Genomes Project database (http://www.internationalgenome.org), as well as by filtering rearrangements identified across an internal panel of normal genomes. TDP status was ascertained as previously described (Menghi et al., 2018).

### RNAseq

RNA-seq libraries for the COH TNBC cohort were generated using the KAPA Stranded mRNA-Seq kit (Roche) according to manufacturer’s instructions and were sequenced on a HiSeq4000 Illumina platform to generate 75 bp paired end reads. RNA-seq libraries for the UW OvCa cohort were generated using the KAPA mRNA Hyperprep kit (Roche) according to manufacturer’s instructions and were sequenced on a NovaSeq Illumina platform to generate 100 bp paired end reads. For both datasets, high quality unique paired-end reads were aligned to the GRCh38-based UCSC gene reference transcriptome using Bowtie2, and RSEM (Li and Dewey, 2011) was used to estimate the abundance of each individual gene. Upper quartile normalization was performed within each tumor sample after discarding genes with no counts. Finally, gene expression levels were adjusted using a percentile rank transformation. Analysis of differential gene expression was performed using the *limma* package in R. For the COH TNBC dataset, TNBC transcriptional subtypes were determined using the TNBCtype-4 tool as described in (Lehmann et al., 2016).

### Published tumor datasets

- *AOCS cohort.* This cohort comprises 80 primary serous ovarian carcinomas that are part of the Australian Ovarian Cancer Study (AOCS). Their *BRCA1/2* status had been previously published (Patch et al., 2015). Structural variant calls were downloaded from the COSMIC data portal (data freeze version v78). Clinical data, RNAseq-based gene expression and methylation array (Illumina human methylation450K array) data were downloaded from the ICGC data Portal (https://dcc.icgc.org/) in August 2020.
- *TCGA OvCa cohort.* This dataset comprises 314 primary serous ovarian carcinomas from the TCGA ovarian cohort (Cancer Genome Atlas Research, 2011). Their BRCA*1/2* mutational status was obtained from cBioPortal (June 2020 download, (Cerami et al., 2012; Gao et al., 2013)) and from previous publications (Cancer Genome Atlas Research, 2011; Maxwell et al., 2017). *BRCA1* promoter methylation was computed based on Illumina human methylation27K array Beta-values downloaded from the UCSC Xena Browser in June 2020: the average Beta-value relative to the four probes mapping to the minimal promoter of the *BRCA1* gene (cg19531713, cg19088651, cg08993267, cg04658354) was computed and a threshold of 0.4 was selected to identify methylated cancers. Clinical data and microarray-based gene expression data were downloaded from the UCSC Xena Browser in June 2020.
- *TCGA TNBC cohort.* A subset of 34 primary TNBCs with unambiguous *BRCA1*meth or *BRCA1*mut status was selected from the TCGA breast cancer cohort (Cancer Genome Atlas, 2012). Clinical data, RNAseq-based gene expression and methylation array data (either Illumina human methylation27K array, n = 15 genomes; or Illumina human methylation450K array, n = 19 genomes) were downloaded from the UCSC Xena Browser in June 2020. *BRCA1* promoter methylation was assessed as described above (*TCGA OvCa cohort*) for cancer genomes analyzed using the 27K arrays. Cancer genomes analyzed using the 450K arrays were classified as methylated when the average of the Beta-values corresponding to the 21 probes mapping to the minimal promoter of the *BRCA1* gene (cg12984107, cg19531713, cg19088651, cg08386886, cg08993267, cg24806953, cg20187250, cg15419295, cg16963062, cg16630982, cg21253966, cg04110421, cg04658354, cg17301289, cg09441966, cg26891576, cg20760063, cg10893007, cg11126247, cg12182452, cg09831010) was higher than 0.25. *Glodzik TNBC cohort.* This dataset comprises 82 TNBC cancer genomes characterized by BRCA1 deficiency (*BRCA1*mut, n = 25; *BRCA1*meth, n = 57), as described by Glodzik et al. (Glodzik et al., 2020). Illumina methylationEPIC array data and RNAseq-based gene expression data were obtained from the Gene Expression Omnibus data repository under accessions IDs GSE148748 and GSE96058, respectively. Beta-values were computed from the methylated and unmethylated probe intensities according to the following formula: β=max(methylated, 0)/(max(methylated, 0)+max(unmethylated, 0)+α), where the constant offset α = 100 is added to regularize the computed Beta-value when both methylated and unmethylated probe intensities are low.

### PDX experiments and response metrics

The main TNBC PDX cohort (n = 33) analyzed in this study were established in the Advanced In Vivo Models Core at the Baylor College of Medicine, under the direction of Prof. Michael T. Lewis (Zhang et al., 2013). Response to single agent carboplatin and docetaxel was assessed as part of a larger ongoing PDX study, details for which to be published in detail elsewhere. Briefly, for each model, appropriately sized mice cohorts were established via orthotopic tumor transplantation and when tumors reached a volume of ∼150-300 mm^3^, mice were randomly assigned to one of three study arms (control, carboplatin or docetaxel) and subjected to a four-week regimen of weekly intraperitoneal doses of either vehicle, carboplatin at 50 milligrams per kilograms of body weight (mg/kg) or docetaxel at 20 mg/kg. Tumor growth was monitored via biweekly tumor volume measurements using calipers, and datapoint collected up to seven days following the administration of the last dose of each agent (i.e., day 28) were used for the response analysis.

Response to treatment was evaluated using two independent metrics that resulted in highly comparable outcomes, as detailed in **Tables S2 and S5**. In the first instance, we took full advantage of the PDX model system and compared tumor growth rates between animals in the treatment and control arms. Briefly, the rate of tumor growth for each animal was computed by fitting a linear curve to the log-transformed tumor volumes, as previously described (Hather et al., 2014). This value was then compared to the average tumor growth rate of the animals in the control arm for each TNBC PDX model, to calculate the percentage difference in tumor growth rate relative to each individual animal. Finally, values corresponding to replicate animals were averaged to compare percentage reductions in tumor growth rates across the different genetic backgrounds. Binary response was determined by setting a threshold corresponding to 5% reduction in tumor growth rate in the treatment arm compared to the control arm. This percentage reduction corresponds to a 50% decrease in tumor volume for the animals in the treatment arm when compared to those in the control arm over a 14-day timeframe, assuming comparable tumor volumes between the two arms at the beginning of the study. Response was also assessed based on the percentage tumor volume change (ΔVol) at day 28 compared with its baseline (day 0). The criteria for response were adapted from Response Evaluation Criteria in Solid Tumors (RECIST) criteria (Eisenhauer et al., 2009) and defined as follows: complete response, Δvol < −80%; partial response, −80% < Δvol < −30%; stable disease, −30% < Δvol < 20%; progressive disease, Δvol > 20%. The mode outcome for each group of replicate animals was reported as the response call for each PDX model examined, with models which demonstrated RECIST values equivalent to complete or partial response being designated as responders.

Cisplatin response relative to the additional *BRCA1* methylated TNBC PDX models represented in Figure 3 and **Table S5** were obtained from the Jackson Laboratory (JAX) PDX Resource, under protocol 12027 approved by The Jackson Laboratory Institutional Animal Care and Use Committee (IACUC) before study initiation. Details regarding the establishment and therapeutic characterization of these models have been previously published (Shultz et al., 2014). The treatment regimen for these models consisted of three weekly doses of cisplatin at 2 mg/kg. Biweekly tumor volume measurements up to seven days following the administration of the last dose (i.e., day 21) were analyzed as described above. Information and data for PDX models from the JAX PDX Resource are publicly available from the PDX Portal hosted by Mouse Tumor Biology Database (MTB; http://tumor.informatics.jax.org/mtbwi/pdxSearch.do) (Krupke et al., 2017).

Four TNBC PDX models from the JAX PDX Resource were additionally analyzed to assess the effect of *in vivo* cisplatin treatment on *BRCA1* expression and promoter methylation. For each model, appropriately sized mice cohorts were established via subcutaneous tumor injection and when tumors reached a volume of ∼150-300 mm^3^, mice were randomly assigned to one of the two study arms (control vs. cisplatin) and subjected to four weekly doses of either vehicle or cisplatin at 2 mg/kg. The TM00097 PDX model was also assessed for its response to the AC→T regimen, which consisted of three weekly doses of both doxorubicin (2 mg/kg) and cyclophosphamide (40 mg/kg), followed by three weekly doses of docetaxel (6 mg/kg). Following treatment, host mice were given a drug holiday until their tumors reached ∼2000 mm^3^ in volume or, for tumors that shrunk following treatment, until they grew back to at least twice the size of their original pre-treatment volume, at which point they were harvested and assessed for *BRCA1* expression via qPCR and promoter methylation via MSP. All animal procedures employed for this additional study were approved by The Jackson Laboratory Institutional Animal Care and Use Committee (IACUC) under protocol number 12027.

### PDX-derived primary cultures

Primary cell cultures were established from cryopreserved fragments of PDX #TM00099, as previously described (Kim et al., 2018). Briefly, tumor fragments were dissociated through incubation with collagenase for one hour, washed and plated on a layer of 3T3-J2 irradiated feeder cells and grown in the culture medium described by Liu et al. (Liu et al., 2012). Following *in vitro* expansion, single cells were isolated via flow cytometry and further expanded to establish individual clonal lines.

### Cell Culture and IC_50_ Determination

The SUM149 and SUM149.B1.S* cell lines were a kind gift by Prof. Chris Lord. SUM149 carries a pathogenic variant of the *BRCA1* gene (c.2288delT, p.N723fsX13) and has lost the wild type form of the gene. Its CRISPR/Cas9-derived daughter clone SUM149.B1.S* harbors an 80-bp deletion downstream of the parental mutation (c.[2288delT;2293del80]), which restores the open reading frame in the parental BRCA1 c.2288delT allele and encodes a functional 1836-amino acid-long BRCA1 protein (Drean et al., 2017). Both cell lines were authenticated by amplification and Sanger sequencing of the *BRCA1* locus harboring the original and secondary mutations and regularly tested for *Mycoplasma* contamination using the MycoAlert PLUS Mycoplasma Detection Kit (Lonza). The two cell lines were maintained in Ham’s F-12 Nutrient Mixture medium with 5% (vol/vol) FBS, 1% (vol/vol) Penicillin-Streptomycin, 0.01 mg/mL bovine insulin and 1 µg/ml hydrocortisone. For IC_50_ value determinations, cells were plated in 96-well plates at a density of 2 × 10^3^ cells per well. After 24 hours in culture, cisplatin (Selleck Chemicals), docetaxel (Selleck Chemicals), doxorubicin (Selleck Chemicals), mafosfamide (Santa Cruz Biotechnology) were added in triplicate wells to the culture medium in two-fold serial dilutions in the range of 0.05 to 205 μM (cisplatin and mafosfamide), 0.05 to 205 nM (docetaxel), or 0.006 to 26 µM (doxorubicin). Cells were incubated for 72 hours before assessing cell viability using a WST-8 assay (Dojindo Molecular Technologies, Inc.). Absorbance values were normalized to control wells (medium only), and IC_50_ values were calculated using the IC_50_ R package (Frommolt and Thomas, 2008). Four independent biological replicate experiments were carried out for each cell line and each treatment, and the Student’s t-test statistic was used to determine the significance of the difference between the average IC_50_ values relative to the two cell lines.

### *BRCA1* promoter methylation analysis

*BRCA1* promoter methylation status for the COH TNBC and PDX datasets was determined by methylation-specific PCR (MSP). Briefly, 100 ng of genomic DNA was subjected to bisulfite conversion using the EpiTect bisulfite kit (QIAGEN). Following clean-up, the converted DNA was used as template for two separate PCR reactions to amplify the proximate *BRCA1* gene promoter region with two sets of primers specific for either unmethylated (forward: 5’-TTG GTT TTT GTG GTA ATG GAA AAG TGT-3’; reverse: 5’-CAA AAA ATC TCA ACA AAC TCA CAC CA-3’) or methylated DNA (forward: 5’-TCG TGG TAA CGG AAA AGC GC-3’; reverse: 5’-AAA TCT CAA CGA ACT CAC GCC G-3’). PCR products were then loaded on a 2% agarose gel and analyzed to visualize the amplification products. Samples with visible amplification of the methylation-specific PCR template were considered methylated. Either DNA from the *BRCA1*-methylated HCC38 TNBC cell line or CpGenome™ Human Methylated and Non-Methylated DNA Standards (Sigma-Aldrich) were included in each bisulfite conversion reaction as controls. *BRCA1* promoter methylation for the UW OvCa dataset was determined by bisulfite conversion using the Zymo Research EZ DNA Methylation-Direct kit followed by MSP, as previously described (Bernards et al., 2018). Methylation specific droplet digital PCR (MS-ddPCR) was carried out to quantify the levels of *BRCA1* promoter methylation in the longitudinal series of PDXs and in the subset of methylated OvCas from the UW cohort, as previously described (Kondrashova et al., 2018).

### BRCA1 gene expression analysis via qPCR

Up to 500 ng of RNA was reverse transcribed using Maxima reverse transcriptase mix (Fermentas). Each qPCR reaction was performed using 15 ng of cDNA as template, 0.5μM forward and reverse primers and SYBR-Green PCR Mix (Fermentas), according to the manufacturer’s instruction and using the ABI7500 system (ABI). Expression of the *BRCA1* gene was evaluated using the following primers: forward: 5’-TTG CAG TGT GGG AGA TCA AG-3’; reverse: 5’-CGC TTC TCA GTG GTG TTC AA-3’. The *SRP14* gene was used as the housekeeping control (forward: 5’-AGG GTA CTG TGG AGG GCT TT-3’; reverse: 5’-GCT GCT GCT TTG GTC TTC TT-3’). Relative *BRCA1* expression was computed using the delta-delta Ct method.

### PCR validation of the *STAT3/NBR2/BRCA1* fusion gene and fusion transcript junctions

To confirm the tandem duplication breakpoint relative to the *STAT3/NBR2* gene fusion observed in the WHIM75 sample, primers were designed to amplify a ∼400 base pair genomic region surrounding the estimated breakpoint junction (forward: 5’-TCT CCT CTC TGG TCC CTT GA-3’; reverse: 5’-TTT TGG TTT CCA ACC AGA GC-3’). PCR products were visualized on a 2% agarose gel and, following PCR purification, sequenced via Sanger sequencing, to identify the exact sequence of the breakpoint junction. To confirm the expression of the *STAT3/BRCA1* fusion transcript, up to 500 ng of RNA was reverse transcribed using Maxima reverse transcriptase mix (Fermentas) and amplified with PCR primers surrounding the estimated fusion junction (forward: 5’-AGC AGC ACC TTC AGG ATG TC-3’; reverse: 5’-GAA GGC CCT TTC TTC TGG TT-3’). PCR products were visualized and sequenced as described above.

### Analysis of differential methylation

We performed an analysis of probe-level differential methylation between *BRCA1*meth and *BRCA1*mut cancers using the *limma* package in R (Ritchie et al., 2015) and the matrix of M-values as the input data (with M-value defined as log2(β/(1−β))). Moderated t-statistics and associated p-values were obtained for each CpG site and significant differences were identified based on FDR-adjusted p-values < 0.05. In addition, we performed an analysis of differential methylation of genomic regions using the *bumphunter* function in the *minfi* R package (Aryee et al., 2014; Jaffe et al., 2012). The analysis was run independently for each one of the five methylation array datasets examined, using methylation Beta-values as the input data, selecting a medium difference cutoff of 0.2 (i.e., 20% difference in Beta-values averages across the two groups) and running 100 permutations to assess the significance of the candidate regions. All other parameters were set as default. Only differentially methylated regions with a family-wise error rate < 0.05 were considered significant and selected for display in Figure S3B.

### Analysis of differential gene expression

Differentially expressed genes between *BRCA1*meth and *BRCA1*mut cancers were identified using the *limma* package in R (Ritchie et al., 2015). Only differentially expressed genes with an FDR-adjusted p-value < 0.05 were considered significant and selected for display in Figure S3C.

### Analysis of immune cell infiltrates

CIBERSORT is a deconvolution algorithm that allows to characterize the immune infiltrate cell composition of complex tissues, including tumor tissues, from their bulk gene expression profiles (Chen et al., 2018). We applied the CIBERSORT relative analytical tool from the Alizadeh lab (https://cibersort.stanford.edu) and the default LM22 signature matrix to quantify 22 different human hematopoietic cell types across our four expression datasets. For RNAseq-based gene expression datasets, we measured gene abundance in Transcripts Per Kilobase Million and disabled quantile normalization, as recommended. We then compared CIBERSORT-generated abundance scores across responders and non -responders using a logistic regression model for datasets with binary response outcomes (i.e., COH TCGA, UW OvCa and AOCS datasets). For datasets with long-term follow up data (i.e., UW OvCa, AOCS and TCGA OvCa datasets), we performed survival analyses using Cox proportional hazard models and splitting the datasets in two based on the median value of the CIBERSORT score of the immune cell type of interest (i.e., M1 macrophages).

## QUANTIFICATION AND STATISTICAL ANALYSIS

Unless otherwise stated, statistical analyses were performed, and graphics produced using the R statistical programming language version 3.6.1 (www.cran.r-project.org). Logistic regression analyses to assess the association between therapy response as a binary outcome and clinical/genetic variables were performed using the *glm* function in R, specifying a binomial distribution and utilizing *logit* as the link function. In survival analyses, the *coxph function* (in *survival* R package) was applied to compute the Cox proportional hazards regression model, using the *Surv* function to create the survival object. All other statistical tests employed are specified in the appropriate Result sections and corresponding figure legends.

## SUPPLEMENTAL TABLES

**Table S1. Cancer cohorts.**

**Table S2. Cancer sample manifest.**

**Table S3. Associations between clinical and genetic variables and chemotherapy response.**

**Table S4. IC50 values for the SUM149 and SUM149.B1.S* isogenic cell lines.**

**Table S5. PDX models of BRCA1 methylated TNBC.**

**Table S6. Methylation array and gene expression datasets for BRCA1meth vs. BRCA1mut analyses.**

---

**Figure S1.**
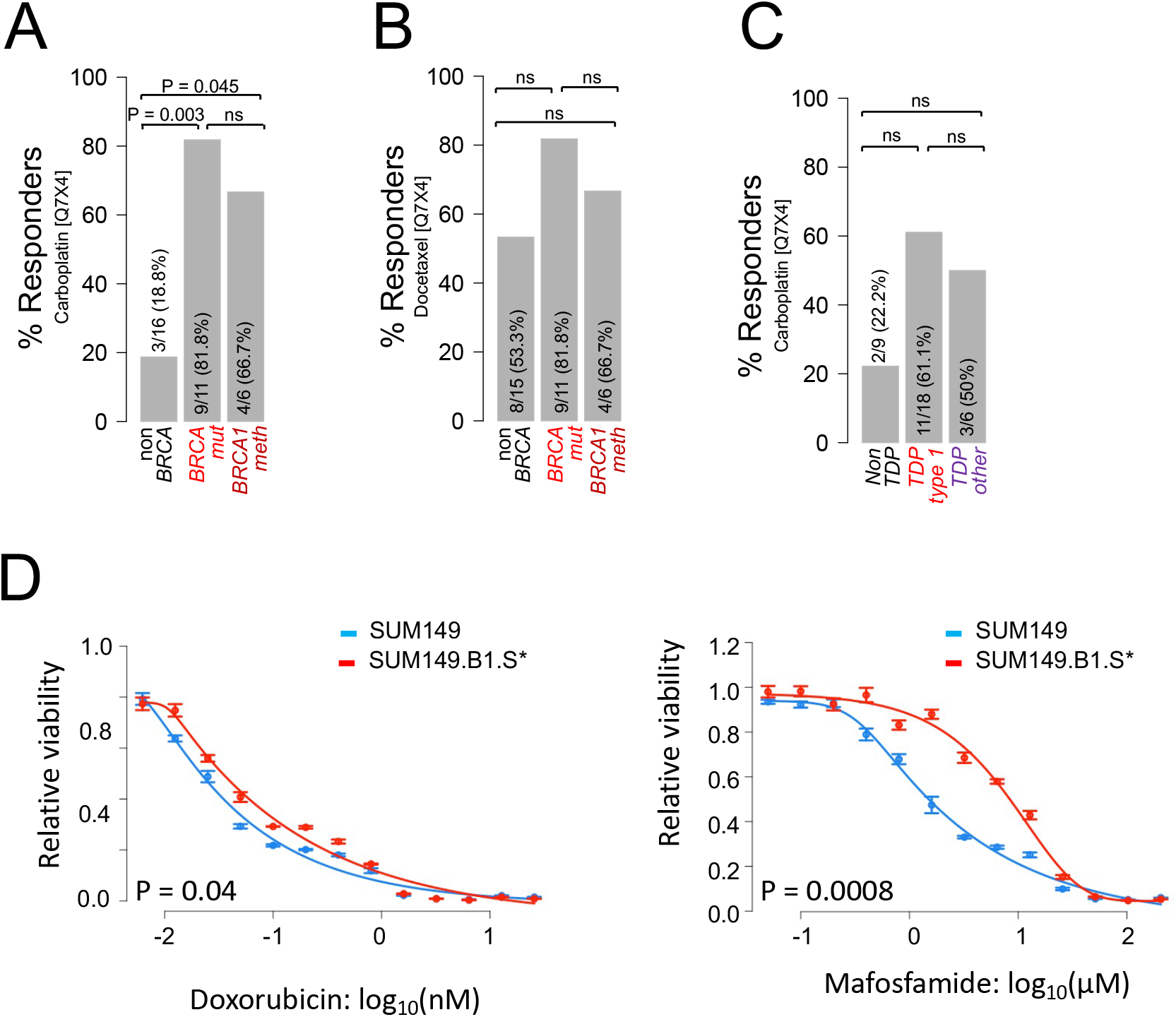
Response to chemotherapy as a function of *BRCA* and TDP status in TNBC PDX models and cancer cell lines. **A-B)** Percentage of carboplatin (**A**) and docetaxel (**B**) responders as a function of *BRCA* status. Responders are defined as animals that achieve complete or partial response as assessed by RECIST criteria. P by logistic regression; ns, not significant. **E)** Percentage of carboplatin responders as a function of TDP status in the PDX cohort. Responders are defined as in (**A-B)**. P by logistic regression. **D)** Doxorubicin and mafosfamide IC50 curves relative to the SUM149/SUM149.B1.S* isogenic cell lines. One representative example of four biological replicate experiments is shown in each graph. Data are presented as mean values and standard errors of the technical replicates. Significance values were calculated using the Student’s t-test (two-tailed) to compare IC50 values from the four biological replicates across the two cell lines.

**Figure S2.**
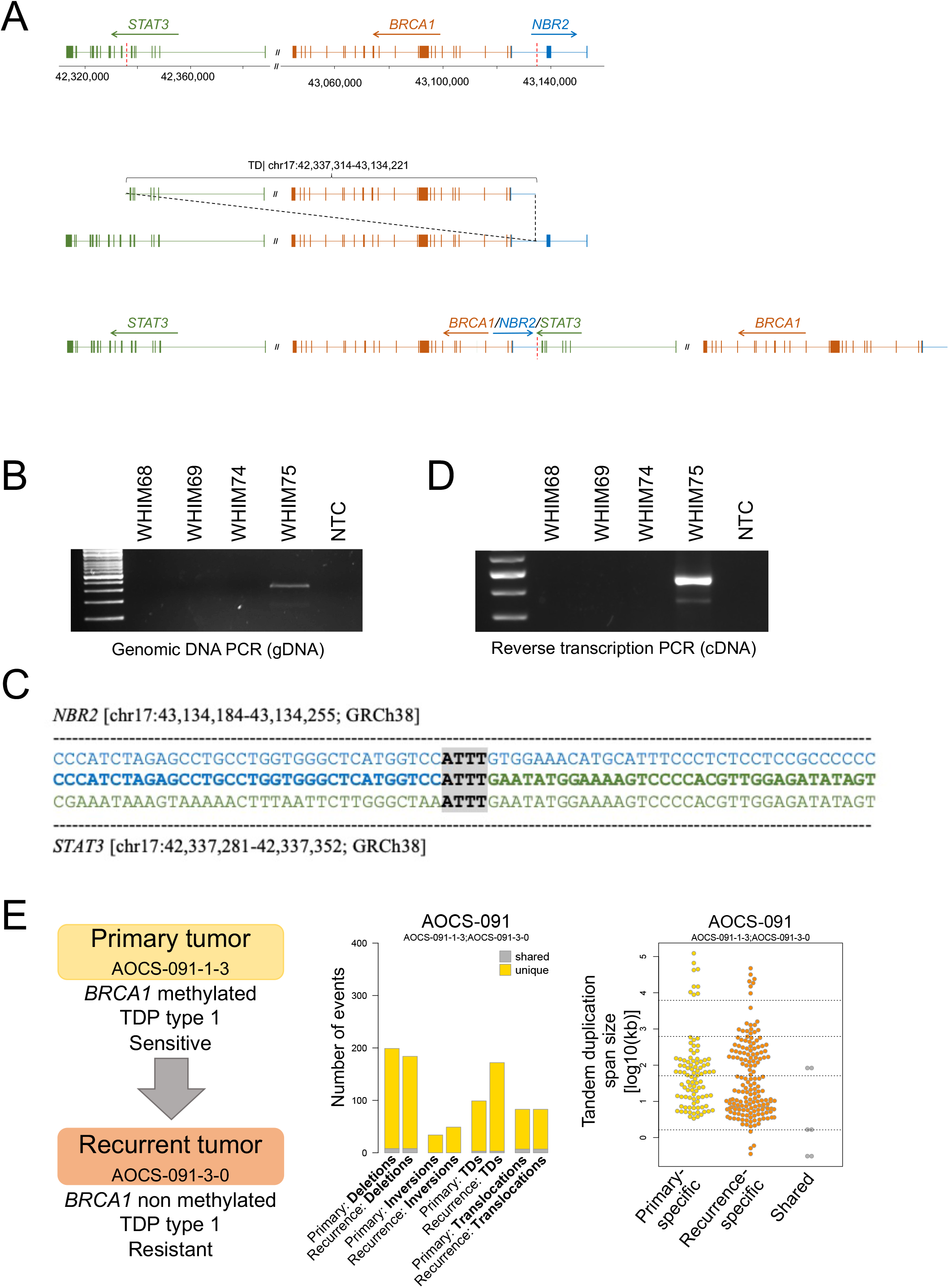
*G*enomic features of acquired resistance to platinum-based therapy in two cases of *BRCA1*meth cancers. **A)** Overview of the genomic region on chr17 hosting the *STAT3, NBR2* and *BRCA1* genes (top), of the de novo tandem duplication in the WHIM75 cancer genome (middle), and of the resulting gene fusion between *STAT3* and *NBR2* (bottom). Individual exons are represented by vertical lines and tandem duplication breakpoint are indicated by red dashed lines. Genomic coordinates are based on the GRCh38 genome built. **B)** Gel image of the genomic DNA PCR using primers spanning the *STAT3/NBR2* gene fusion junction. Only WHIM75 shows an amplification signal. **C)** Sequence analysis of the *STAT3/NBR2* tandem duplication breakpoint junction in the WHIM75 PDX. Sequence alignments relative to the *NBR2* and *STAT3* intronic regions involved in the fusion are shown in blue and green, respectively. The novel gene fusion sequence is shown in the middle row in bold highlight. Grey highlights represent the region of microhomology at the breakpoint junction. **D)** Gel image of the reverse transcription PCR using primers spanning the *STAT3/BRCA1* fusion transcript junction. Only WHIM75 shows an amplification signal. **E)** Summary of structural variations and tandem duplication span sizes relative to the paired primary and recurrent OvCa genomes from donor AOCS-091 (AOCS cohort).

**Figure S3.**
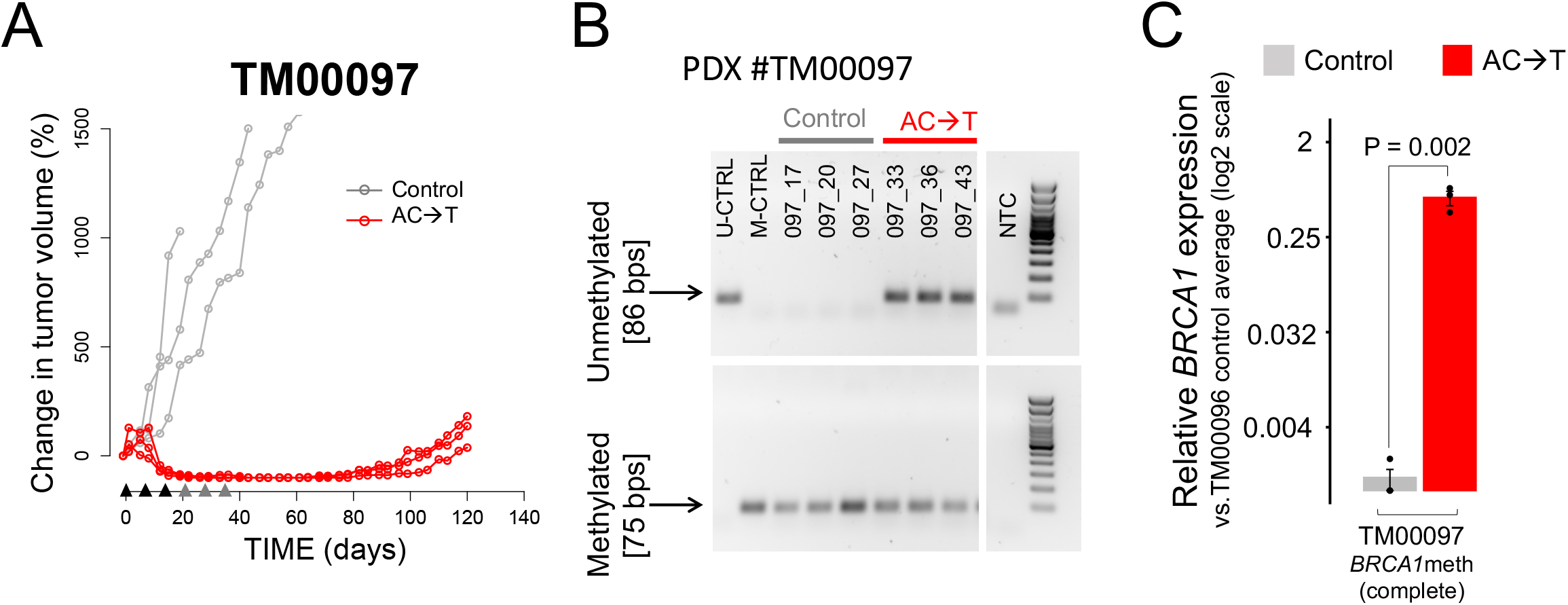
Assessment of *BRCA1* methylation and expression in a *BRCA1*meth TNBC PDX treated with AC→T *in vivo*. **A)** Tumor growth curves relative to TNBC PDX #TM00097. Black and grey arrow heads indicate the timing of the three weekly doses of the doxorubicin/cyclophosphamide combination and of docetaxel, respectively. **B)** MSP results for vehicle tumors and AC→T-treated tumor recurrences relative to PDX #TM00097. **C)** qPCR of *BRCA1* gene expression across vehicle tumors and AC→T-treated tumor recurrences. BRCA1 expression levels are normalized to the average *BRCA1* expression of control tumors from the non*BRCA* PDX #TM00096. P value by Student’s t-test (two-tailed).

**Figure S4.**
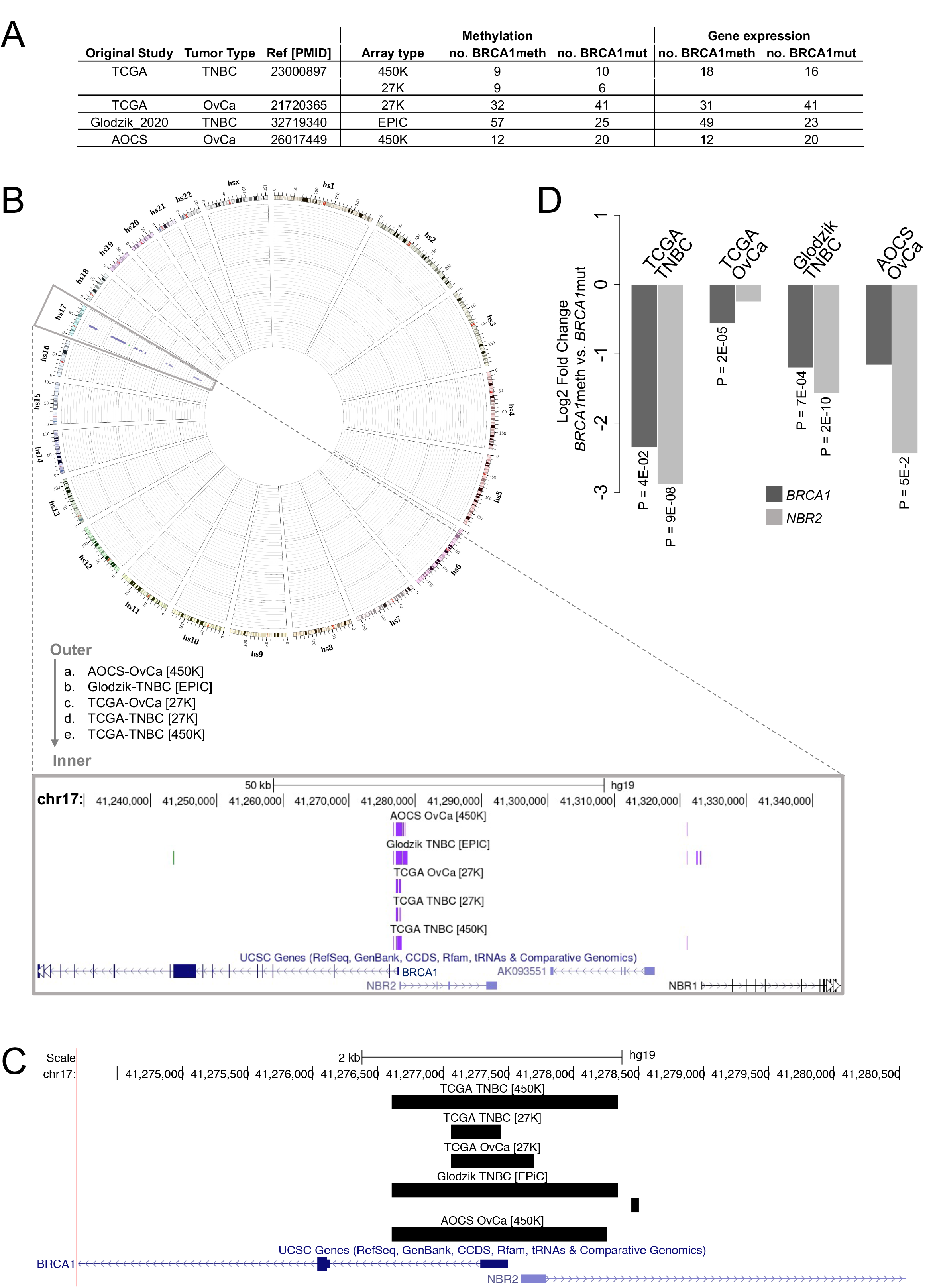
Differential DNA methylation and gene expression between *BRCA1*meth and *BRCA1*mut cancer genomes. **A)** Description of the datasets included in the analyses of differential methylation and differential gene expression between *BRCA1*meth and *BRCA1*mut cancers. **B**) Circos plot of fold changes in methylation values (M-values) between *BRCA1*meth and *BRCA1*mut cancers for each one of the five methylation array datasets examined. Each track spans the −2 to 7 fold-change range, with horizontal lines drawn at each incremental unit. Only significantly differentially methylated CpG sites are depicted (adjusted p-values < 0.05), with purple and green representing CpGs with higher and lower methylation values in the *BRCA1*meth cancer subgroup, respectively. All the significantly differentially methylated CpGs are located at, or in proximity of, the *BRCA1* promoter region. A zoomed-in view of the significant CpG locations is provided as a UCSC Genome Browser screenshot. **C)** UCSC Genome Browser screenshot of the differentially methylated genomic regions (dmrs) between *BRCA1*meth and *BRCA1*mut cancers across the five methylation array datasets. Dmrs were identified using the *bumphunter* function in the *minfi* R package. Analysis of each dataset identified one (4/5 datasets) or two (1/5 datasets) significant dmrs, all of which mapped to the *BRCA1/NBR2* minimal promoter region. **(D)** Bar plot of log2 fold changes in gene expression between *BRCA1*meth and *BRCA1*mut cancers across four independent datasets. Analysis of each dataset identified either *BRCA1* or *NBR2* or both as the only differentially expressed genes between the two groups. Adjusted p-values are shown only for the significant comparisons (FDR < 0.05).

**Figure S5.**
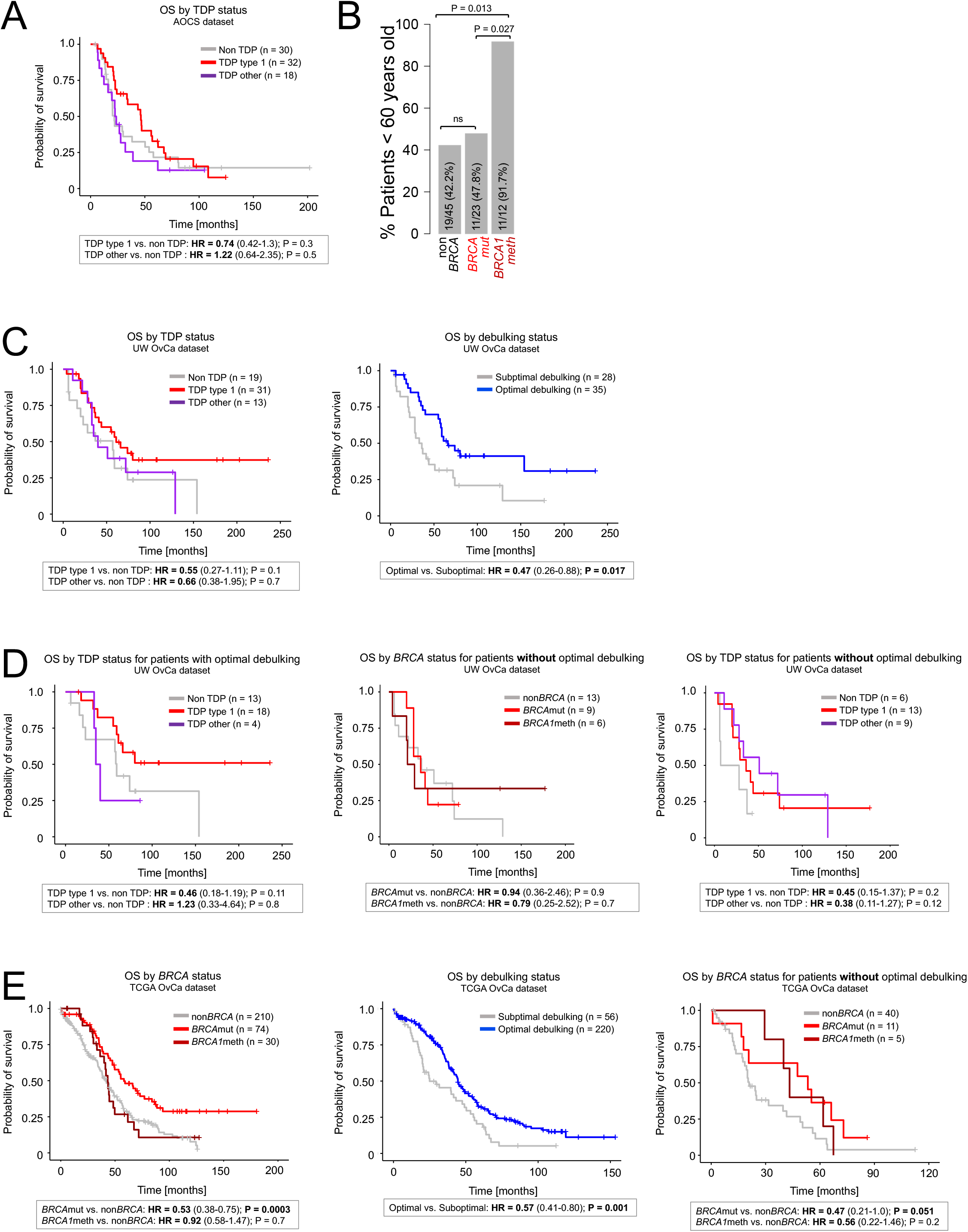
Analysis of therapeutic response in the OvCa cohorts. **A)** AOCS cohort. Overall survival (OS) of patients stratified based on TDP status. **B)** AOCS cohort. Patient age distribution relative to *BRCA* status; P value by logistic regression. **C)** UW OvCa cohort. Overall survival of patients stratified based on based on TDP status (left) and debulking status (right). **D)** UW OvCa cohort. Survival analysis restricted to patients that either received (left) or did not receive optimal debulking (middle and right), stratified based on TDP status or *BRCA* status. **E)** TCGA OvCa cohort. Overall survival of patients stratified based on *BRCA* status (left), debulking status (middle) and restricted to the subset of patients who did not receive optimal debulking (right). For all the survival analyses, the Cox proportional hazards regression model was used to compute hazard ratios (HR) with 95% confidence intervals (in brackets), and their corresponding P values.

**Figure S6.**
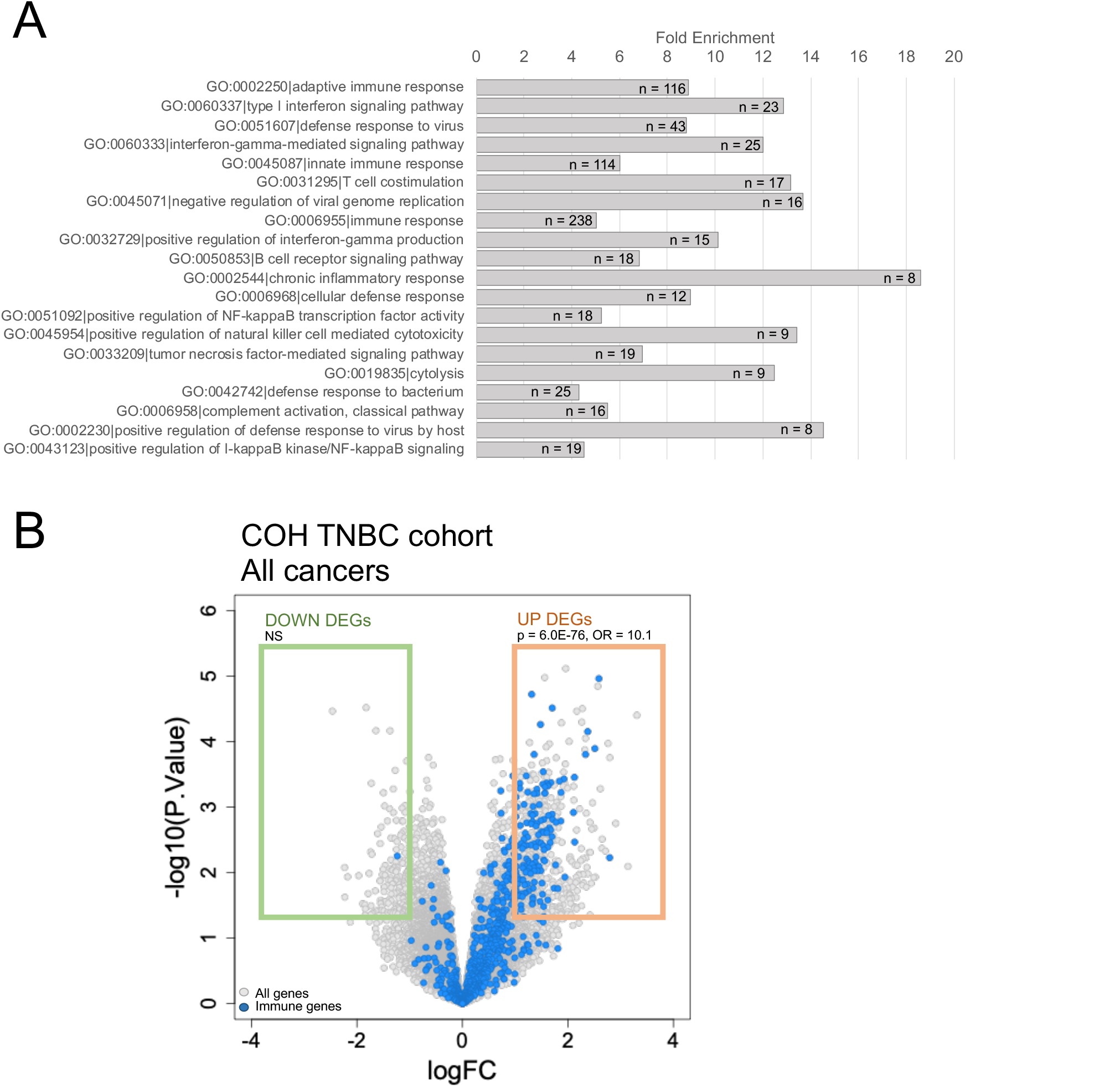
Analysis of transcriptional profiles associated with pCR in the COH TNBC cohort. **A)** TOP 20 Gene Ontology biological process terms significantly enriched in the set of up-regulated genes in TNBCs from patients who achieved pCR vs. those who did not. Terms are sorted by increasing p-value and their relative fold enrichment is indicated on the x axis. The number of significant genes for each category is depicted within each bar. **B)** Volcano plot of differential gene expression between patients who achieved pCR vs. those who did not within the COH TNBC cohort. Significantly differentially expressed genes (p-value <0.05 and absolute log2 fold change > 1), are indicated by green (DOWN) and orange (UP) boxes. Immune genes are shown in blue. Enrichment of immune genes within the up-regulated differentially expressed genes was computed by Fisher’s exact test.

**Figure S7.**
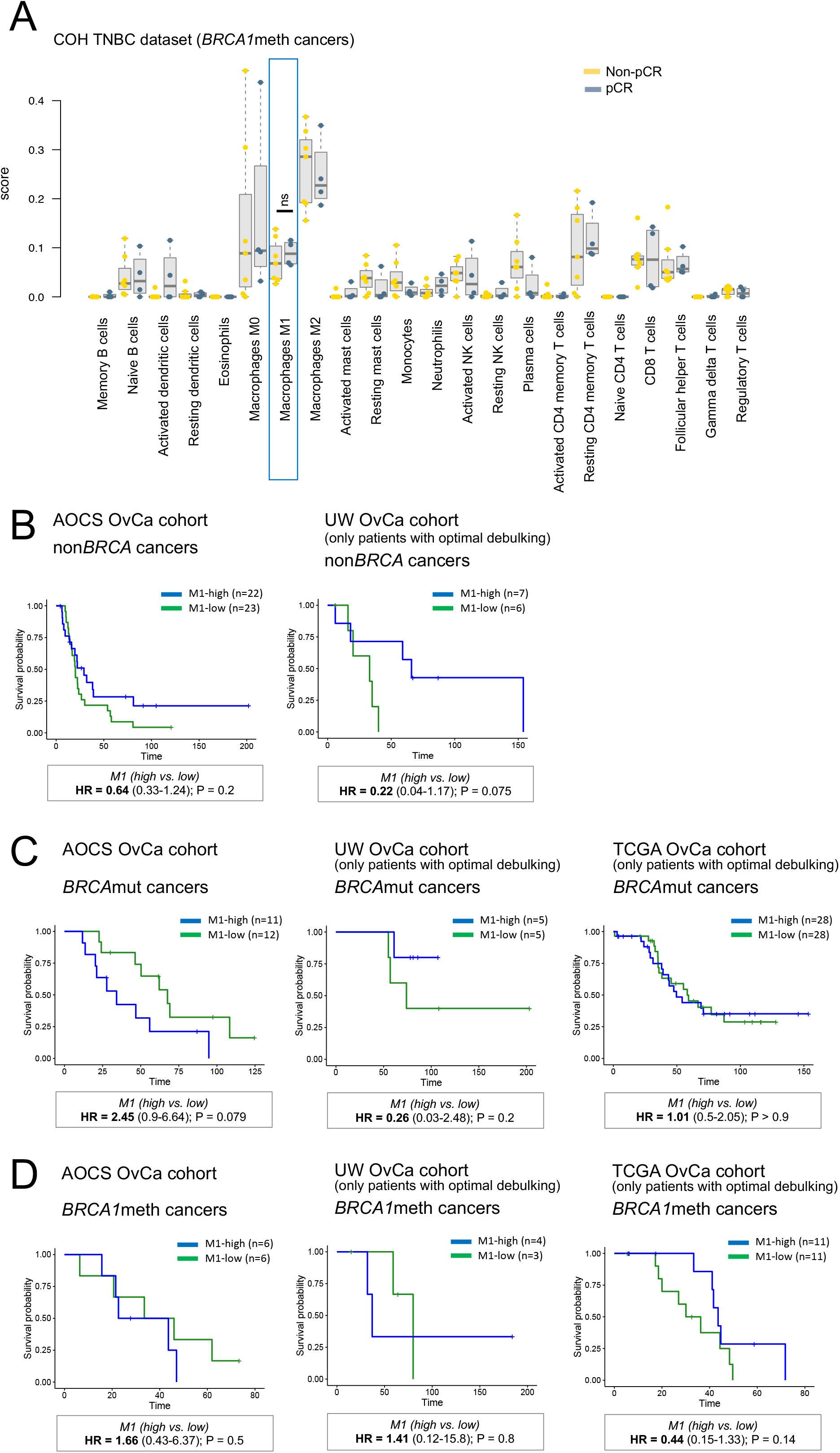
CIBERSORT immune cell type scores across TNBC and OvCa cohorts. **A)** Comparison of CIBERSORT immune cell type scores between tumors from pCR vs. non-pCR patients, in the subset of the COH TNBC cohort with *BRCA1*meth cancers. None of the comparisons are significant (Mann-Whitney test). **B)** Survival analysis for the TCGA OvCa patient cohort (left, all cancers; right, non*BRCA* cancers only), with patients stratified based on their M1 macrophage CIBERSORT score (median split). **B-D)** Overall survival for OvCa patients with non*BRCA* cancers **(B),** *BRCA*mut cancers **(C)** or *BRCA1*meth cancers **(D)**, and high or low levels of the CIBERSORT-derived M1 macrophage signature scores (i.e., M1-high and M1-low, split on the median level for each indicated group). HR, hazard ratio; CI, confidence interval; P value by Cox proportional hazards regression model.

